# DeepPrep: An accelerated, scalable, and robust pipeline for neuroimaging preprocessing empowered by deep learning

**DOI:** 10.1101/2024.03.06.581108

**Authors:** Jianxun Ren, Ning An, Cong Lin, Youjia Zhang, Zhenyu Sun, Wei Zhang, Shiyi Li, Ning Guo, Weigang Cui, Qingyu Hu, Weiwei Wang, Xuehai Wu, Yinyan Wang, Tao Jiang, Theodore D. Satterthwaite, Danhong Wang, Hesheng Liu

## Abstract

Neuroimaging has entered the era of big data. However, the advancement of preprocessing pipelines falls behind the rapid expansion of data volume, causing significant computational challenges. Here, we present DeepPrep, a pipeline empowered by deep learning and workflow manager. Evaluated on over 55,000 scans, DeepPrep demonstrates a 11-fold acceleration, exceptional scalability, and robustness compared to the current state-of-the-art pipeline, providing a promising solution to meet the scalability requirements of neuroimaging.

## Main Text

Advances and transparency in neuroimaging are propelled by the rapidly growing amounts of publicaly available data from large-scale projects (e.g., the UK BioBank (UKBB) with 50,000+ scans^1^), data sharing initiatives (e.g., the openneuro^2^), and international consortiums (e.g., the ENGIMA^3^) (see Extended Data Fig. 1). Neuroimaging data typically requires complex and multi-stage preprocessing pipelines to enable accurate brain tissue segmentation, spatial normalization, and other essential preprocessing steps. While prevailing preprocessing pipelines such as FreeSurfer^4^, fMRIPrep^5^, QSIPrep^6^, and ASLPrep^7^ have led to numerous important findings, their original design for relatively small sample sizes makes them hard to meet the scalability demands in the era of big data. Concurrently, clinical applications of neuroimaging, such as imaging-guided neuromodulation^8^, require fast turn-around time and robustness at the individual level. This becomes particularly challenging when dealing with patients who exhibit brain distortions induced by traumas, gliomas, or strokes^9,10^. To fulfill these emerging requirements, a computationally efficient, scalable, and robust preprocessing pipeline is needed.

Hence, we propose DeepPrep, a highly efficient, scalable, and robust preprocessing pipeline for neuroimaging powered by deep learning algorithms and a workflow manager. Deep learning-based algorithms for neuroimaging have shown great potential in improving computational efficiency and robustness^11-14^. Workflow managers are widely used to manage complex workflows for big data processing in various fields, such as bioinformatics, due to their scalability, portablibity, and computational resource efficiency^15^. However, this has not been fully exploited in the field of neuroimaging. Here, we applied DeepPrep to structural and functional MRI. We integrated multiple learning-based modules, including FastCSR^13^, SUGAR^12^, SynthMorph^14^, and FastSurferCNN^11^ (Fig. 1a, Supplementary Table 1 for all dependencies). These deep-learning modules facilitate computational efficiency in typically time-consuming operations, such as cortical surface reconstruction, surface registration, anatomical segmentation, and volumetric spatial normalization. All software modules are linked by 83 discrete yet interdependent task processes, which are packaged into a Docker or Singularity container along with all dependencies^16^. Our incorporation of Nextflow -- a reproducible, scalable, and portable workflow manager^15,17^ -- enables our pipeline to maximize computational resources utilization through dynamically scheduling parallelization. The workflow manager also makes it convenient to deploy the pipeline in diverse computing environments, including the local computers, high-performance computing (HPC), and cloud computing (Fig. 1b). Importantly, DeepPrep is a BIDS-App that can automatically configure appropriate preprocessing workflows based on the metadata stored in the brain imaging data structure (BIDS) layout^18^. In addition to providing preprocessed structural and functional data (see Supplementary Fig. 1 for an example), DeepPrep also generates a visual report for each participant and a summary report for a group of participants to facilitate data quality assessments by adapting from MRIQC^19^ (Supplementary Fig. 2), as well as a detailed runtime report (Supplementary Fig. 3). Documentation (https://deepprep.readthedocs.io/en/latest/) and code base (https://github.com/indilab/deepprep) of DeepPrep is version controlled and actively updated, enabling users and developers to participate in the further development of the pipeline.

**Fig. 1.**
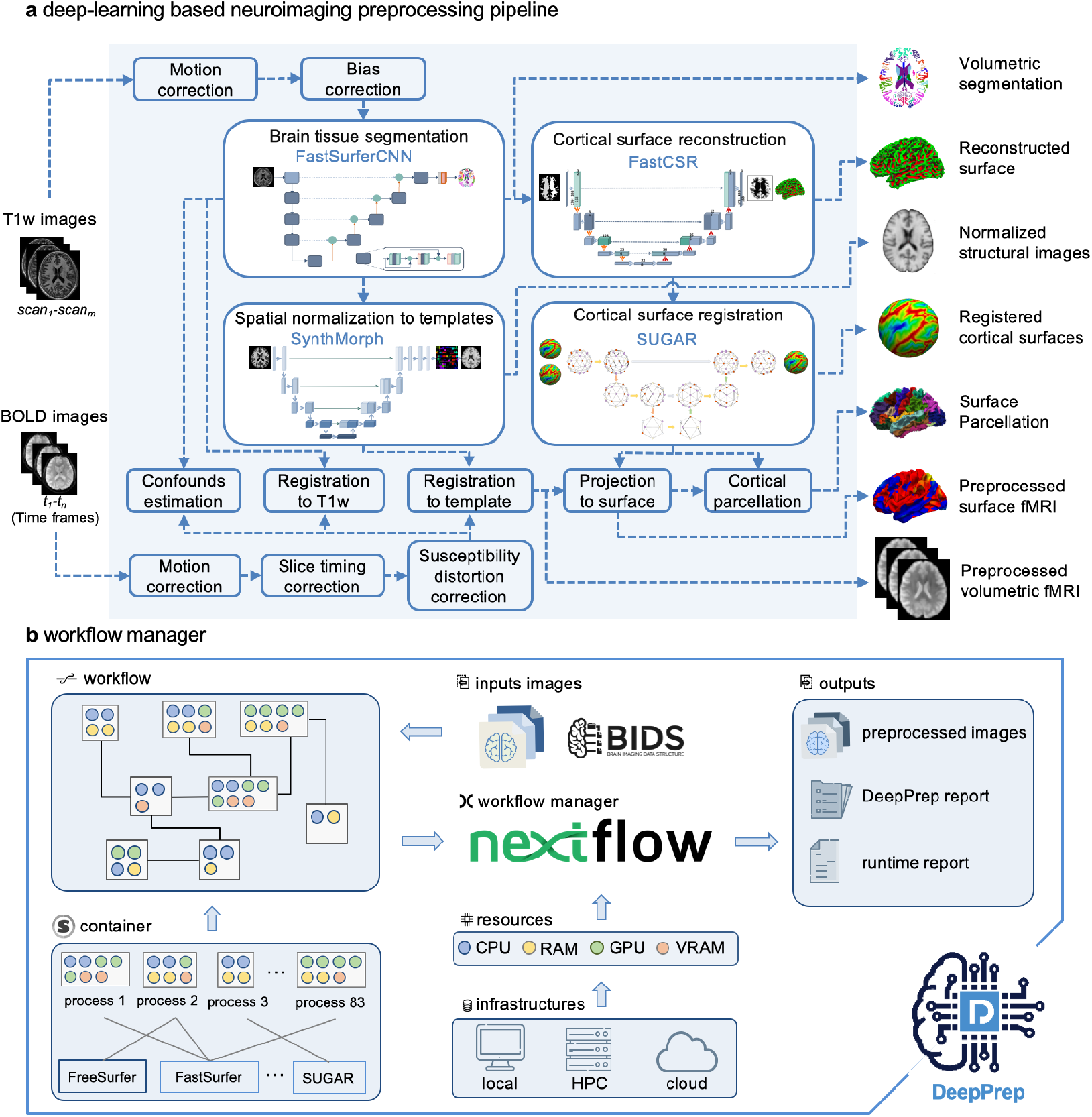
A computationally efficient and scalable neuroimaging pipeline is empowered by deep-learning algorithms and workflow managers. **a**, The neuroimaging pipeline leverages deep learning algorithms, including FastSurfer, FastCSR, SUGAR, and SynthMorph, to replace the most time-intensive modules present in conventional pipelines. This substitution enables the achievement of highly efficient and robust brain tissue segmentation, cortical surface reconstruction, cortical surface registration, and volumetric spatial normalization. The current version of the pipeline supports both anatomical and functional MRI preprocessing in both volumetric and cortical surface spaces. **b**, The preprocessing pipeline is organized into multiple relatively independent yet interdependent processes. Imaging data adhering to the BIDS format are preprocessed through a structured workflow managed by Nextflow, an open-source workflow manager designed for life sciences^17^. Nextflow efficiently schedules task processes and allocates computational resources across diverse infrastructures, encompassing local computers, HPC clusters, and cloud computing environments. The pipeline yields standard preprocessed imaging and derivative files, DeepPrep quality control reports, and runtime reports as its outputs.

To demonstrate the performance of DeepPrep, we applied it to over 55,000 scans from 8 datasets acquired with diverse populations, scanners, and imaging parameters (Supplementary Table 2). These datasets include structural and functional MRI data from the UKB^1^ for evaluating computational efficiency and scalability, one manually labelled brain dataset (the Mindboggle-101 dataset^20^) for evaluating brain segmentation, two precision neuroimaging datasets (the MSC dataset ^21^ and the CoRR-HNU dataset ^22^) for accuracy and reliability assessment, and three clinical datasets (the CRRC-Stroke dataset, SHH-DoC dataset, and BTH-Glioma dataset) for robustness assessment. These data were also processed using the current state-of-the-art processing pipeline, fMRIPrep^5^, for comparison.

In a local workstation equipped with CPUs and a GPU (see Methods), DeepPrep successfully preprocess 1,189 subjects from the UKB dataset per week, with an average processing time of 8.5 minutes per subject. This processing speed represents approximately 11-fold faster than that achieved by fMRIPrep in the same workstation, which processed 107 subjects with an average processing time of 92.6 minutes per subject (Fig. 2a & b). Moreover, when we performed separate preprocessing for anatomical and functional scans, DeepPrep exhibited acceleration factors of 13.8 and 12.1 times, respectively (Extended Data Fig. 2). Notably, in a HPC environment, the trade-off between preprocessing time and computational resource utilization, measured in terms of CPU hours and associated costs, becomes a critical consideration. Achieving shorter preprocessing time through recruiting more CPUs comes with higher expense on hardware (Fig. 2c), which can be a significant concern for users. To explore this trade-off relationship, we conducted an analysis by recruiting one, two, four, eight, and sixteen CPUs for processing. FMRIPrep exhibited a characteristic curve, illustrating the trade-off between processing time and costs (Fig. 2c). In contrast, DeepPrep demonstrated stability in both processing time and associated costs, due to its computational flexibility in dynamically allocating computational resources to match the specific requirements of individual task processes. Importantly, the computational costs associated with DeepPrep were found to be at least 7.5 times lower than those of fMRIPrep. Additionally, we demonstrated the scalability of DeepPrep by successfully preprocessing the entire UKB neuroimaging dataset, consisting of over 54,515 scans, within 6.5 days in a HPC cluster (see Methods).

**Fig. 2.**
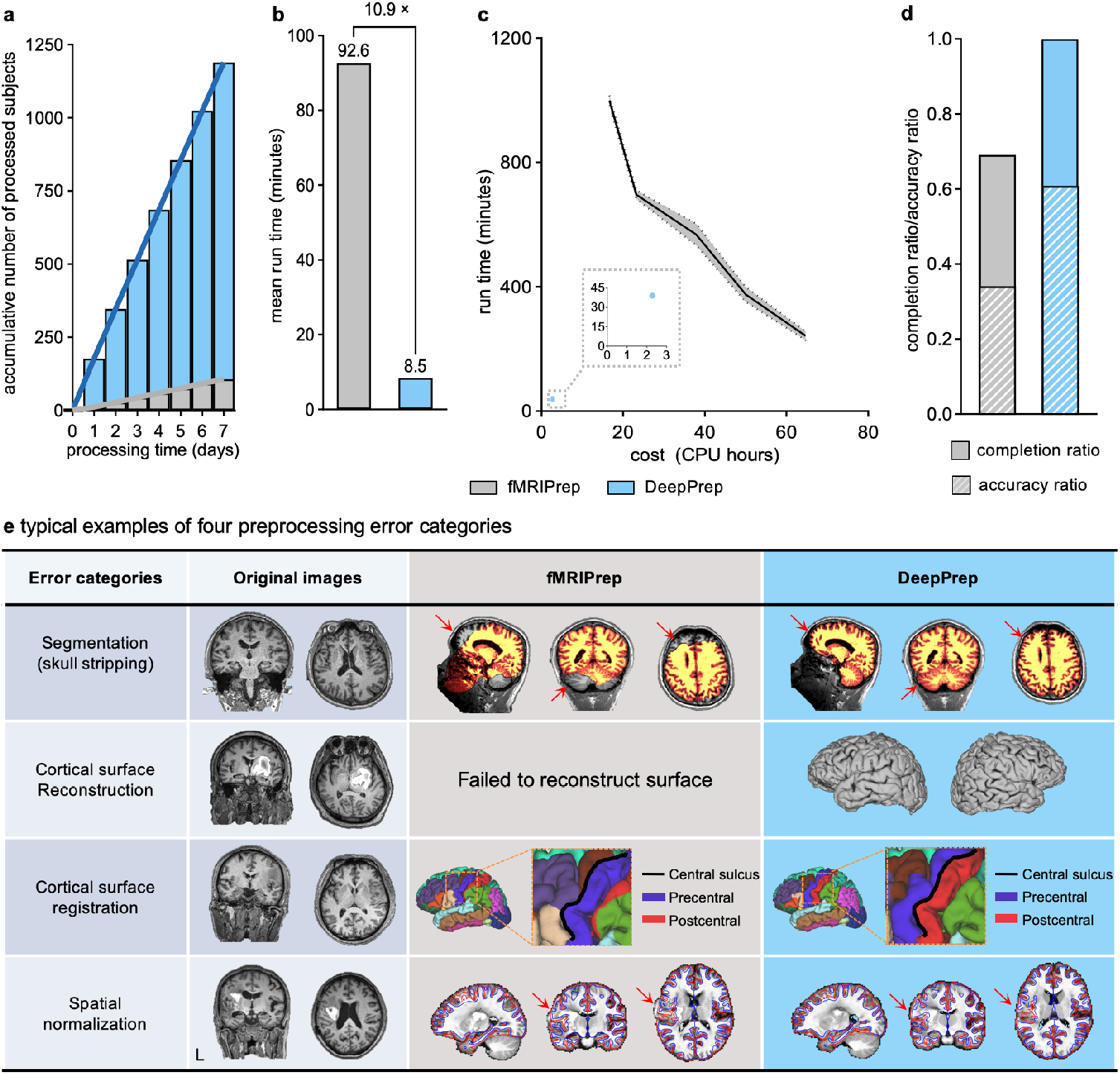
DeepPrep achieves over 10-fold acceleration and shows robustness in processing clinical samples. **a**, DeepPrep (blue bars) successfully processed 1189 participants’ scans from the UKB dataset on a workstation, whereas fMRIPrep (gray bars) processed 107 participants. **b**, Average processing time per participant of DeepPrep is 8.5 minutes, with a remarkable 11-fold increase compared to fMRIPrep’s 92.6 minutes. **c**, In the HPC context, preprocessing time of fMRIPrep can be reduced by allocating additional computational resources for processing each single scan, yet with higher hardware expense associated with CPU hours (the gray line). DeepPrep offers flexibility in resource allocation, tailoring computational resources to the specific requirements of each task process, resulting in reliable costs and efficient processing time for individual participants (the blue dot). The cost of DeepPrep is at least 20 times lower than that of fMRIPrep. **d**, Robustness of DeepPrep is assessed in preprocessing intractable clinical samples. DeepPrep successfully completed preprocessing in 100% of patients, with 60.4% of patients being correctly preprocessed. Meanwhile, fMRIPrep’s success rate was 69.8% for completion and 34.0% for correct preprocessing. **e**, Preprocessing errors are categorized into four types, including brain tissue segmentation, cortical surface reconstruction, cortical surface registration, and volumetric spatial normalization. Four representative cases with preprocessing errors are presented, illustrating obvious brain lesions or imaging noises in the original images. fMRIPrep yields an inaccurate brain mask when skull stripping, failed to reconstruct cortical surfaces, exhibited misalignment in surface parcellation in the pre- and post-central gyrus, and produced inappropriate volumetric normalization in perilesional region. In contrast, DeepPrep successfully and accurately processed these cases.

The DeepPrep and fMRIPrep outputs were also compared in the Mindboggle-101, MSC, and CoRR-HNU datasets, covering a range of aspects including anatomical parcellation and segmentation (Extended Data Fig. 3), anatomical morphometrics (Extended Data Fig. 4), spatial normalization (Extended Data Fig. 5), temporal signal-to-noise ratio (Extended Data Fig. 6), task-evoked responses (Extended Data Fig. 7), functional connectivity (Extended Data Fig. 8), and test-retest reliability in functional connectomes (Extended Data Fig. 9). Overall, the comparison indicated that DeepPrep consistently yielded preprocessing results that were either highly similar or even superior to those generated by fMRIPrep when assessed using various metrics. These results support the accuracy of DeepPrep while concurrently maximizing efficiency.

Next, we screened 53 clinical samples that could not be successfully processed by FreeSurfer v6.0 within 48 CPU hours due to distorted brain or imaging noises from three clinical datasets (see Methods). These samples were preprocessed using both DeepPrep and fMRIPrep. The results indicated that DeepPrep exhibited a higher pipeline completion ratio (100.0%), a higher preprocessing accuracy ratio (60.4%), and significantly shorter processing time for each subject (45.5 ± 5.6 minutes) compared to fMRIPrep (Fig. 2d; completion ratio: 69.8%, chi-square test: χ^2^ = 16.6, *p* = 4.7×10^−5^; accuracy ratio: 34.0%, χ ^2^ = 7.4, *p* = 0.006; processing time = 513.2 ± 108.4 minutes, two-tailed paired t-test, *t*_(52)_ = -26.2, *p* < 0.0001). The occurrences of fMRIPrep preprocessing failures and errors could be attributed to four main causes: segmentation errors, surface reconstruction failures, surface registration errors, and volumetric registration errors (Fig. 2e). Intriguingly, these four causes correspond to four processing modules where deep-learning algorithms replace conventional algorithms, indicating the robustness of DeepPrep in handling complicated clinical cases.

In summary, DeepPrep pipeline demonstrates exceptional efficiency and robustness in processing large sample neuroimaging datasets and complex clinical cases. This success can be attributed to the integration of workflow managers for handling big data and the utilization of deep learning algorithms. With time, we plan to expand DeepPrep into a comprehensive platform for processing multimodal neuroimaging. While the current version focuses primarily on the most time-consuming workflows in structural and functional MRI, future versions will integrate additional modalities, such as arterial spin labeling (ASL) and diffusion imaging (dMRI). These enhancements will benefit the broader neuroimaging community.

## Methods

DeepPrep offers a computationally efficient, robust, and scalable neuroimaging preprocessing pipeline for investigators, incorporating recent advancements in deep learning for neuroimaging processing and dynamic workflow management. Designed to be compatible with the BIDS standard^18^, DeepPrep facilitates automatic workflow configurations with minimal manual intervention, utilizing metadata provided by BIDS. Moreover, DeepPrep is facilitated by container and workflow manager technologies, and thus is computationally efficient and portable across different computing environments, providing a highly efficient alternative for neuroimaging preprocessing. For optimial acceleration, it is highly recommended to excute DeepPrep in a computing eviornment equipped with GPUs, although it remains executable on CPU-only systems (see Supplementary Note 3 for minimum system requirements). Additionally, version control of the project’s codes and documents is managed through GitHub. While we plan to expand DeepPrep to be a comprehensive pipeline for multimodal neuroimaging in the future, the current version primarily focuses on the most time-consuming workflows, including anatomical preprocessing, functional preprocessing, and imaging quality control. To evaluate its computational efficiency, scalability, accuracy, and robustness, we preprocessed over 55,000 scans from seven datasets using DeepPrep and compared the results with fMRIPrep^5^ and FreeSurfer^4^.

### Preprocessing workflow for anatomical images

#### Overall Description

The anatomical preprocessing workflow in DeepPrep closely follows the FreeSurfer pipeline while efficiently replacing several of the most time-consuming steps with deep learning algorithms. Specifically, the volumetric anatomical segmentation, cortical surface reconstruction, and cortical surface registration are accomplished using FastSurferCNN^4^, FastCSR^13^, SUGAR^12^, respectively. Detailed information on the training and test procedures for each algorithm is provided in Supplementary Note 2. The remaining steps in the workflow remain consistent with FreeSurfer v7.2, including the spherical mapping, morphometric estimation, and statistics. These steps ensure the continued reliability and accuracy of the overall preprocessing process while harnessing the benefits of deep learning algorithms to enhance computational efficiency. The preprocessing workflow consists of several essential steps as follows:

#### Motion correction

If multiple T1w images are available for each participant or each session, FreeSurfer’s “recon-all -motioncor” is employed to correct head motions across the scans. This process yields an average T1w image to minimize the impact of head motions on data quality.

#### Segmentations

The whole brain is segmented into 95 cortical and subcortical regions using FastSurferCNN^11^. Specifically, the segmentation model utilized is FastSurferCNN, which is optimized for accurate and rapid anatomical segmentations.

#### Skull-stripping and bias field correction

A brain mask is generated according the 95 whole-brain regions to achieve accurate and robust skull-stripping. The T1w images undergo N4 bias field correction using SimpleITK with reference to a brain mask. Afterward, the normalized and skull-stripped T1w images could be fed into the subsequent steps.

#### Cortical surface reconstruction

The white-matter and pial cortical surfaces are reconstructed based on the anatomical segmentation derived from the FastSurferCNN^11^. This process utilizes FastCSR^13^, a deep-learning-based model designed to accelerate cortical surface reconstruction. FastCSR leverages an implicit representation of the cortical surface through the level-set representation, and uses a topology-preserving surface extraction method to yield white and pial surfaces represented by triangulated meshes.

#### Cortical surface registration

The reconstructed surface is inflated to a sphere with minimal distortion using the FreeSurfer command *mris_sphere*. Both rigid and non-rigid registrations for the spherical cortical surface are performed to align anatomical landmarks and morphometrics, facilitated by the deep-learning cortical surface registration framework, SUGAR^12^. Individual spherical surfaces are aligned to the fsaverage template surfaces by default.

#### Cortical surface parcellation

The cortical surface parcellation is generated based on the cortical surface registration using the FreeSurfer command *recon-all -cortparc*. Subsequently, the cortical parcellation is projected to the volumetric segmentation by assigning voxels their closest cortical labels via the command *mri_surf2volseg*, thereby replacing the cortical parcellation derived from FastSurferCNN.

### Preprocessing workflow for functional images

#### Overall Description

The functional preprocessing workflow in DeepPrep incorporates advanced registration methods, SynthMorph^14^, to replace the most time-consuming step, the spatial normalization. The workflow is also complemented by modules from existing tools, including AFNI^23^, FSL^24^, and fMRIPrep^5^, to form a comprehensive functional image preprocessing method. The fMRI preprocessing workflow consists of several essential steps as follows:

#### Motion correction and slice-timing correction

The head motion parameters of the BOLD fMRI signals are estimated by MCFLIRT from FSL^25^, with the reference volume generated from averaging non-steady state volumes or dymmy scans. Slice-timing correction is included in our processing pipeline for fMRI data using 3dTshift from AFNI^23^, when slice-timing information is available in the BIDS metadata. This is an optional step and can be deactivated if the BIDS metadata does not specify slice times.

#### Susceptibility distortion correction

DeepPrep incorporates SDCFlows (Susceptibility Distortion Correction Workflows)^26^ to correct susceptibility distortions. SDCFlows offers versatile workflows designed to preprocess various MRI schemes, enabling the estimation of B0 field-inhomogeneity maps directly associated with distortion. This distortion correction is applied to the fMRI data when the appropriate fieldmap information is available within the BIDS metadata. Distortion correction is an optional step.

#### Coregistration

A rigid registration is performed using FreeSurfer’s boundary-based registration to align motion-corrected fMRI volumes to native T1w images for each subject. The registration optimizes the boundary-based loss function to align the boundary between gray and white matter across different imaging modalities.

#### Spatial normalization

The spatial normalization step aims to normalize individual brain images to a standard template, such as the MNI152NLin6Asym volumetric template and FreeSurfer’s fsaverage6 surface template. Traditionally, this step is highly time-consuming due to the requirement of non-rigid surface and volumetric registration, often taking hours for computations. However, in DeepPrep, a deep-learning algorithm called SynthMorph^14^ is utilized to achieve both rigid and non-rigid volumetric registration in minutes. Additionally, for both rigid and non-rigid cortical surface registration, SUGAR is used to achieve accurate alignment in anantomical landmakrs and morphometrics in seconds. Subsequently, preprocessed BOLD fMRI volumes are projected to the MNI152NLin6Asym template and fsaverage6 template surfaces by default, through applying deformation matrices derived from the registrations. The pipeline also flexibly supports normalization to other volumetric human brain templates managed by the TemplateFlow _27_.

In summary, this preprocessing workflow utilizes a combination of conventional methods and advanced deep learning algorithms to efficiently and accurately preprocess structural and functional images for neuroimaging analysis.

### The workflow manager and containers

In the DeepPrep, we used the Nextflow^17^ (https://www.nextflow.io), an open-source workflow manager widely used in life science, and used Docker (https://www.docker.com) and Singularity^16^ containers to establish a scalable, portable, and reproducible neuroimaging preprocessing pipeline across diverse computing environments.

Nextflow facilitates the DeepPrep in optimizing computational resource utilization, particularly in HPC clusters and cloud infrastructures, and ensuring cross-platform portability and reproducibility. Specifically, DeepPrep established and executed the pipeline using key Nextflow components, including process, channel, workflow, and executor, with some modifications aimed at improving the GPU utilization efficiency. Initially, we disassembled the pipeline into 83 discreate minimal executable steps, enabling parallelization and substantial improvements in computational efficiency. The disassemble also allowed for precise monitoring and control of system resource consumption for each individual step. Each step was defined as a Nextflow process, ensuring that input and output data adhered to expected standards and specifying the system resource requirements for each process. Data flow pipelines were delineated using the Nextflow channel component, and the complete workflow was specified through the Nextflow workflow component, integrating predefined processes and data flow pipelines.

Once the pipeline was established, DeepPrep employed the Nextflow executor component, serving as an abstraction layer between the processes and the underlying execution system. This abstraction facilitated the portable execution of the pipeline across diverse platforms and infrastructures. However, in cases where the execution system was set to ‘local’, the existing version of Nextflow lacked GPU resource monitoring capabilities. Given the benifits in computational efficiency of DeepPrep from multiple GPU-intensive processes, concurrent execution on a single GPU-equipped machine could result in errors due to insufficient GPU video memory (VRAM). To address this challenge, we devised a GPU scheduling module, utilizing two distributed mutual exclusion locks based on Redis (https://redis.io). This module categorized GPU processes into two groups: those consuming less than half of the VRAM and those requiring more than half. Processes with lower VRAM requirements could proceed upon acquiring a single lock, while those with higher demands needed to secure both locks simultaneously. This module effectively fixed GPU-related errors and substantially improved GPU utilization efficiency before Nextflow’s GPU monitoring implementation.

In addition, we prioritized the reproducibility of our preprocessing pipeline. To this end, DeepPrep underwent rigorous version control and was fully compatible with execution in both Docker and Singularity containers. Singularity, designed for scientific computational tasks such as neuroimaging processing, offered the flexibility to execute across a spectrum of computing infrastructures and platforms, ranging from local machines to HPC clusters and cloud environments. Notably, Singularity emphasized security, a paramount concern in scientific computing, particularly in HPC settings. It operated within a user-space model, obviating the need for elevated privileges, thereby substantially reducing security risks typically associated with superuser access.

### DeepPrep report

DeepPrep automatically generates a descriptive HTML report for each participant and session (Supplementary Fig. 2). The report commences with a concise summary of key imaging parameters extracted from the BIDS meta information. Subsequently, the report provides an overview of the overall CPU and GPU processing times for the data preprocessing. Key processing steps and results for structural images are visually presented, including segmentation, parcellation, spatial normalization, and coregistration. The normalization and coregistration outcomes are demonstrated through dynamic ‘before’ versus ‘after’ animations. Additionally, the report includes a carpet plot, showcasing both the raw and preprocessed fMRI data, along with a temporal signal-to-noise ratio (tSNR) map. Finally, the report concludes with comprehensive boilerplate methods text, offering a clear and consistent description of all preprocessing steps employed, accompanied by appropriate citations.

### Standard output

The preprocessed structural MRI data are organized to align with the results of FreeSurfer, encompassing the normalized and skull-stripped brain, reconstructed cortical surfaces and morphometrics, volumetric segmentation, cortical surface parcellation, and their corresponding statistics. Additionally, transformation files for surface spherical registration are included, as illustrated in Supplementary Fig. 1, depicting the data structure. For the preprocessed functional MRI data, the naming adheres to the BIDS specification for derived data. The default output spaces for the preprocessed functional MRI consist of three options: 1. the native BOLD fMRI space, 2. the MNI162NLin6Asym space, and 3. the fsaverage6 surfaces space. However, users have the flexibility to specify other output spaces, including the native T1w space and various volumetric and surface templates available on TemplateFlow. The main outputs of the preprocessed data include:

1. Preprocessed fMRI data
2. Reference volume for motion correction
3. Brain masks in the BOLD native space, include the nuisance masks, such as the ventricle and white-matter masks.
4. Transformation files for between T1w and the fMRI reference and between T1w and the standard templates.
5. Head motion parameters and the temporal SNR map
6. Confound matrix

### Evaluation data

#### Data Description

We collected over 55,000 scans from 8 datasets to evaluate the performance of DeepPrep. The evaluation datasets are distinct from training and validation sets for deep learning models to avoid information leakage. The training and validation sets for these models are described in previous reports^11-14^. To achieve comprehensive evaluations, we collected over 55,000 scans from seven datasets as test sets. A large-scale dataset (the UKBB dataset^1^, 49300 participants with 5215 participants including repeated scans, totally 54515 anatomical and functional scans) was used to evaluate the computational efficiency, a dataset with manual annotations of the anatomical regions (the Mindboggle-101 dataset^20^, n = 101) was used to evaluate anatomical parcellation, and two repeated measured datasets [the Midnight scanning club (MSC) dataset^21^ (10 participants and 10 sessions per participant) and CoRR-HNU dataset^22^ (30 participants and 10 sessions per participant)] were used to evaluate the accuracy and test-retest reliability. Moreover, to evaluate the robustness in preprocessing clinical cases with anatomical distortions, we included three in-house patient datasets, including the Glioma dataset, the DoC dataset and the Stroke dataset. See the *Supplementary Note 1* for more detailed description of each dataset.

### Comparison to alternative preprocessing tools

We compared the processing time and preprocessed anatomical and functional outcomes from DeepPrep with those from fMRIPrep (version: 22.0.2) and FreeSurfer (version: 7.2.0), respectively.

#### Processing time and computational costs

The preprocessing time of both the DeepPrep and fMRIPrep was recorded for comparison and evaluated in the identical hardware environment (a workstation with an Intel Core i9 10980XE 3.00GHz × 36 CPU and a NVIDIA RTX3090 GPU with 24 GB RAM). To examine the scalability of the DeepPrep in large-scale datasets, we also preprocessed 54515 scans from 49300 participants in the UKBB dataset using our pipeline in an HPC cluster of CPL. The cluster is equipped with 1920 CPU cores (2.90GHz) and 20 NVIDIA RTX3090 GPUs. To quantify the computational costs in the HPC environment, we calculated CPU hours, derived by multiplying the number of recruited CPU cores by the duration of usage. Of note, due to the reliance on GPU processing for DeepPrep, we empirically converted one GPU hour into an equivalent of 15 CPU hours to enable a standardized comparison, according to the cloud computing charge^1,2^.

#### Assessments of structural imaging preprocessing

Structural images were preprocessed using both the DeepPrep and fMRIPrep. The preprocessed structural results were compared between the two pipelines in anatomical segmentation and parcellation and similarities in the morphometrics, as elaborated below.

#### Accuracy of anatomical parcellation and segmentation

The surface-based parcellation and volumetric segmentation of structural images were automatically delineated by different pipelines. We first assessed the similarity, measured by Dice coefficient, between the parcellations and segmentations automatically generated by different preprocessing pipelines and manually delineated annotations considered as the ‘ground truth’. The higher Dice coefficient indicates greater similarity and better accuracy. To visually demonstrate the differences in parcellation and segmentations, we directly estimated the difference percentages in Dice coefficients from two pipelines.

#### Similarity to the morphometric atlas

To assess the surface registration performance, we compared the similarities in the morphometrics, including sulcal depth and curvature, between the aligned surfaces and the FreeSurfer ‘fsaverage’ template surface. We employed the mean squared error (MSE) to measure the dissimilarity, with lower MAE values indicating better alignment registration performance. We used the metrics to assess registration performance because the primary objective of surface registration is to align these anatomical features between individual cortical surfaces and atlases^12,28,29^.

#### Assessments of functional imaging preprocessing

Preprocessed functional images were compared between the DeepPrep and fMRIPrep in multiple aspects, including spatial normalization, tSNR, task activations, and functional connectivity. To fairly compare, we performed identical post-processing pipelines for task and resting-state fMRI, controlling preprocessed images to the same smoothing level. For the task fMRI, we used the FSL FLIM^30^ (FMRIB’s Improved Linear Model) to perform a standard general linear model for the first level analysis after high-pass temporal filtering (100s). The second level inference was performed using the FLAME^31^ (FMRIB’s Local Analysis of Mixed Effects), based on the first-level outcomes. For post-processing resting-state fMRI, we included the bandpass filtering (0.01-0.08Hz) and regression of nuisance variables, including 6 motion parameters, white-matter signal, ventricular signal, whole-brain signal, and their temporal derivatives.

#### Spatial normalization

After normalizing the BOLD-fMRI images to the MNI152NLin6Asym volumetric template, we calculated the standard deviation map of averaged BOLD time series across participants, in order to examine the performance of spatial normalization. A higher standard deviation around the brain outline indicates a lower performance of spatial normalization. ***tSNR***. The voxel-wise tSNR was calculated to assess the amount of informative signal relative to the noise level in preprocessed images, following a previous report ^32^. The tSNR was estimated by the TSNR of Nipype, which first applied a quadratic detrending, then calculated the temporal mean and divided by its temporal standard deviation for each voxel of each participant. The individual voxel-wise tSNR map was averaged across all participants.

#### Motor-task activation map

The MSC motor task^21^ is a block design with conditions of tongue, left hand, right hand, left foot, right foot, adapted from the paradigm used in the Human Connectome Project^33^. Task-evoked activations were modeled by a general linear model (GLM) for each voxel and each session for the first-level analyses. The second level analyses averaged data across session for each participant based on the fixed effect model. The third level analyses grouped data across participants based on the mixed effect model. The task post-processing analyses were performed using the FSL^24^.

#### Seed-based cortico-cortical and cortico-cerebellar functional connectivity

To assess the similarity in resting-state functional connectivity (RSFC) derived from different pipelines, we examined typical long-range RSFC of a cortico-cortical and a cortico-cerebellar circuit. Specifically, we identified cortical and cerebellar seeds from previous studies ^34^. For the cortico-cortical circuit, a seed in left-hemisphere posterior cingulate cortex (PCC; MNI coordinate = -2, - 53, 26) was selected. For the cortico-cerebellar circuit, a seed in the left angular gyrus (AG, MNI coordinate = -49, -63, 45) was selected. Each seed region of interest (ROI) was a 6-mm spheres centered around the MNI coordinates of each seed. The RSFC was estimated by Pearson’s correlation coefficient between average BOLD time series within each seed ROI. To normalize the distribution of correlation coefficients, these r values were converted to z values through the Fisher’s r-to-z transformation. To demonstrate both the surface-based and volumetric RSFC, we performed surface-based analyses for the PCC seed and volumetric analyses for the AG seed. The similarity in RSFC was assessed in the MSC dataset.

#### Test-retest reliability in functional connectomes

To assess test-retest reliability in functional connectomes, we utilized a volumetric atlas comprising 300 cortical and subcortical ROIs ^35^ and a cortical surface atlas comprising 300 cortical ROIs^36^ to estimate the whole-brain and cortical functional connectomes, respectively. The test-retest reliability was measured by calculating the similarity between functional connectomes obtained from two different segments in the repeated measured datasets, namely, the MSC dataset and the CoRR-HNU dataset. Furthermore, we investigated the dependence of reliability on scanning duration by randomly selecting and concatenating two or more sessions to measure the functional connectome and its corresponding test-retest reliability, following the procedure described previously^21^.

### Application to data in clinical setting

To assess the pipelines’ robustness in handling clinical cases with distorted brains, we included a total of 53 scans from three clinical datasets, comprising patients with various conditions, including stroke, glioma, and disorders of consciousness (see Supplementary Note 1 for more details). Notably, the selected structural images of these clinical cases either failed or yielded significant errors during original preprocessing using the conventional FreeSurfer v6.0 procedure. The criterion for processing failure is that the procedure cannot be completed within 48 hours on a CPU. To compare the pipelines’ performance, we employed both DeepPrep and fMRIPrep to preprocess all cases and recorded the processing time, completion and accuracy rates, as well as analyzed the error reasons. Completion is defined as successfully producing the processed outputs within 48 hours on 8 CPUs and one GPU.

#### Processing success and accuracy rate

The completion rate was determined as the proportion of clinical datasets that successfully completed all preprocessing steps, resulting in intact output files. On the other hand, the accuracy rate was defined as the proportion of samples whose preprocessing results were deemed accurate, considering the rationality of surface reconstruction, anatomical segmentation, registration quality, and functional connectivity. The assessment of processing accuracy was conducted independently by three experienced neuroimaging experts through visual inspection (J.R., W.C., and H.L.). The experts were blinded to the label of the preprocessing pipeline to minimize subjective biases. A sample was classified as accurate when at least two neuroscientists reached a consensus on its accuracy.

#### Preprocessing error categories

During the assessment of failed and inaccurate samples, we conducted a systematic step-by-step inspection of the preprocessing outputs to identify the underlying reasons for the errors. Based on this analysis, the failures were categorized into four distinct groups: segmentation errors, surface reconstruction failures, surface registration errors, and volumetric registration errors. Segmentation errors were observed in cases where clear inaccuracies were present in the segmentation of anatomical tissues. These errors were more commonly found in samples with extensive lesions or substantial head motion during T1w imaging scanning. Surface reconstruction failures were characterized by either the failure to reconstruct cortical surfaces or an excessively prolonged processing time (exceeding 48 hours) to produce results. These issues often arose due to challenges in accurately fixing surface topology in distorted brains. Both surface and volumetric registration errors indicated misalignment of morphometric features, such as sulci and gyri. Such misalignments could lead to inaccuracies in the registration of brain structures.

### Meta-analysis of large-scale neuroimaging studies

To summarize recent trends in big data of neuroimaging, we conducted a meta-analysis of large-scale neuroimaging studies. First, we performed a PubMed (http://www.pubmed.gov) search on studies published before June 30, 2023, using the keywords: (large scale OR large sample size) AND (neuroimage OR *MRI). Next, we collected further studies by reviewing the reference lists of relevant papers, including studies that employed open-source big data or reviews focusing on large-sample size studies. The inclusion criteria for large-scale datasets were as follows: (1) the study recruited a sample larger than 1000 healthy participants or 400 participants from specific populations, such as patients, twins, or preterm brains; (2) the study was peer-reviewed; and (3) the study contained MRI data with at least one type of structural and functional images. These literature searches yielded a total of 42 studies, comprising over 200,000 participants. Each sample was then categorized into one of three types of projects based on its funding resources. The first type was named “Big Project” as it was supported by a single funding source. The second type was named “Consortium” as it involved international collaboration with multiple funding sources. The last type was named “Data Sharing Platform” as it shared data on the same or different topics from different groups.

### Statistical Analysis

Chi-square tests were used for comparisons in completion and accuracy ratios. Processing time, Dice coefficients of cortical parcellation and subcortical segmentations, similarities in functional connectomes of two pipelines were statistically compared using a two-tailed paired-sample t-test. False Discovery Rate (FDR) correction was used to account for multiple testing.

## Acknowledgements

This work was supported by Changping Laboratory (2021B-01-01), China Postdoctoral Science Foundation (2022M720529, J.R.; 2023M730175, W.C.), Beijing Natural Science Foundation (JQ23040), and Natural Science Foundation of China (82072786).

## Footnotes

1 https://aws.amazon.com/cn/ec2/instance-types/r7g/

2 https://aws.amazon.com/cn/ec2/instance-types/g5/

