## Supplementary Fig. 1 for "DeepPrep: An accelerated, scalable, and robust pipeline for neuroimaging preprocessing empowered by deep learning"

**Supplementary Information**

Supplementary Notes


Supplementary Note 1: Dataset description

Supplementary Note 2: Deep-learning algorithm description

Supplementary Note 3: Minimum system requirements

Supplementary Tables

Supplementary Table.1 | software and toolbox dependencies

Supplementary Table.2 | Characteristics of evaluation datasets

Supplementary Table.3 | Comparisons of anatomical segmentation in subcortical structures between fMRIPrep and DeepPrep

Supplementary Table.4 | Comparisons of anatomical segmentation in cortical regions between fMRIPrep and DeepPrep

Extended Figures

Supplementary Figures

Supplementary Fig. 1 | An example of output files

Supplementary Fig. 3 | An example of the DeepPrep report

Supplementary Fig. 3 | An example of the runtime report

#### Supplementary Notes

#### Supplementary Note 1: Dataset description

##### *Evaluation datasets*

The performance evaluation of our proposed pipeline involved more than 55,000 participants from eight different datasets. We performed a comprehensive assessment, examining various aspects including computational efficiency, scalability, accuracy, test-retest reliability, and robustness in clinical samples. Detailed descriptions of each dataset can be found below, and the dataset characteristics are summarized in Supplementary Table 2.

***UKB dataset.*** The UKB dataset was utilized to assess computational efficiency and scalability. This extensive dataset comprised 49,300 participants, ranging in age from 44 to 83 years (mean age = 64.37 ± 7.98 years, 25,639 women and 23661 men). Notably, 5,215 participants underwent scans on two separate visits, resulting in a total of 54,515 pairs of structural and functional MRI data. Data acquisition was performed using 3T Siemens Skyra scanners equipped with 32-channel head coils. Each participant underwent the acquisition of a three-dimensional T1-weighted structural image at a high resolution of 1 mm isotropic, employing a 3D MPRAGE sequence. FMRI data were obtained using an echo planar imaging (EPI) sequence (2.4 × 2.4 × 2.4 $mm^3$, TR = 0.735 s, multiband acceleration factor of 8, 490 timepoints, 6 minutes 10 seconds). During the fMRI acquisition, participants were instructed to maintain their eyes open and focus on a central crosshair. The UK Biobank ensured full consent from all participants, and the study received ethical approval from the North West Multicenter Research Ethics Committee. Access to the UKB dataset was granted through access application 99038. Additional details regarding the UK Biobank can be found on their official website at https://www.ukbiobank.ac.uk/.

***MSC dataset.*** The Midnight Scan Club (MSC) dataset serves as a representative precision fMRI dataset, for evaluating preprocessing accuracy and test-retest reliability. This dataset comprises ten participants (5 women, age= 29.1 ± 3.3 years). Each participant underwent ten 30-minute rs-fMRI sessions and ten 10-minute motor task fMRI sessions, resulting in a total of 400 minutes of fMRI data. All imaging data were acquired using a Siemens TRIO 3T MRI scanner. For each participant, T1-weighted structural MRI data (voxel size=0.8×0.8×0.8 $mm^3$, TE=3.74 ms, TR=2400 ms, TI=1000 ms, flip angle=8°, 224 sagittal slices). Functional imaging data, including rs-fMRI, were obtained using a gradient-echo EPI sequence (TR=2.2 s, TE=27 ms, flip angle=90°, voxel size=4 × 4 × 4 $mm^3$, 36 slices). Prior to data acquisition, informed consent was obtained from all participants, and the study received approval from the Washington University School of Medicine Human Studies Committee and Institutional Review Board. Further details about the dataset can be found in previous studies[1].

***CoRR-HNU dataset.*** Another precision fMRI dataset, the CoRR-HNU dataset, was utilized to assess preprocessing accuracy and test-retest reliability. This dataset comprised 30 young, healthy participants (15 women; 20-30 years old). Each participant underwent ten sessions, with each session including a T1w image and a 10-minute resting-state fMRI (rsfMRI) scan. Imaging data were acquired using a GE Discovery MR750 3T MRI scanner equipped with an 8-channel head coil. For T1w structural images, a 3D "spoiled-gradient-echo" sequence was employed (voxel size = 1.0×1.0×1.0 $mm^3$, TR = 8.06 ms, TE = 3.1 ms, flip angle = 8°, 176 sagittal slices). Functional data were acquired through an EPI sequence (TR=2000 ms, TE=30 ms, flip angle=90°, voxel size=3.4×3.4×3.4 $mm^3$, 43 slices). Ethical approval for the study was granted by the Ethics Committee of the Center for Cognition and Brain Disorders at Hangzhou Normal University. Furthermore, written informed consent was obtained from each participant prior to their enrollment in the study. A more detailed description of the dataset could be found in the previous report[2].

***Mindboggle-101 dataset.*** The Mindboggle-101 dataset constitutes a valuable resource consisting of 101 labeled brain images that have undergone manual correction. These labels adhere to a consistent human cortical labeling protocol known as the DKTatlas[3]. This dataset serves as a crucial validation tool for assessing the anatomical parcellation within the preprocessing pipeline. Remarkably, the Mindboggle-101 dataset stands out as the largest and most comprehensive collection of freely accessible, manually labeled human brain images. It includes anatomically labeled brain surfaces and volumes, all derived from T1-weighted images of healthy individuals. A subset of the dataset, which includes five subjects (specifically, MMRR-3T7T-2, Twins-2, and Afterthought-1), was acquired specifically for this dataset. The remaining subjects were sourced from publicly available datasets, such as Test-Retest OASIS1, the Multi-Modal Reproducibility Resource, Nathan Kline Institute Test–Retest, Nathan Kline Institute/Rockland Sample, Human

Language Network subjects, and the Colin Holmes 27 template. A more detailed description of the dataset could be found in the previous report[3].

***Clinical samples***

To evaluate the robustness in preprocessing clinical samples with extremely distorted brains, we collected 53 patients with distorted brains from three clinical datasets. We screened extremely distorted brains through preprocessing them by FreeSurfer v6.0, yielding 53 out of 424 patients who failed to be processed within 48 CPU hours. The remaining samples were used to evaluate the robustness in clinical samples.

***BTH-Glioma dataset.*** We screened 19 out of 168 patients with brain gliomas in Beijing Tiantan Hospital (BTH). Their T1w images were acquired on a Siemens TimTrio 3T scanner with a sagittal 3D T1w sequence (TR = 2530 ms, TE = 3.37 ms, flip angle = 7°, voxel size = 1 × 1 × 1 $mm^3$). RsfMRI data were obtained using an EPI sequence (TR = 2000 ms, TE = 30 ms, flip angle = 90°, voxel size = 3.44 × 3.44 × 3.750 $mm^3$). Each patient underwent an 8-minute refMRI scan. Written informed consent was obtained from each patient in accordance with guidelines and regulations previously approved by the IRB of BTH, Capital Medical University.

***SHH-DoC dataset.*** We screened 15 out of 38 patients with DoC caused by brain trauma in Shanghai Huashan Hospital (SHH). Their T1w images were acquired on a Siemens MAGNETOM 3T scanner with a sagittal 3D T1w sequence (TR = 1000 ms, TE = 2.15 ms, flip angle = 8°, FOV = 256 × 256, 176 sagittal slices, voxel size = 1 × 1 × 1 $mm^3$). RsfMRI data were obtained using an EPI sequence (TR = 2000 ms, TE = 30 ms, flip angle = 90°, voxel size = 3 × 3 × 3 $mm^3$). Five 6-minute rsfMRI runs were acquired for each patient. Written informed consent was obtained from each patient in accordance with guidelines and regulations previously approved by the IRB of Shanghai Huashan Hospital, Fudan University.

***CRRC-Stroke dataset.*** We screened 19 out of 218 patients with DoC caused by brain trauma in Beijing Bo'ai Hospital, China Rehabilitation Research Center (CRRC). Their T1w images were acquired on a Philips Ingenia 3T scanner with a sagittal 3D T1w sequence (TR = 7.13ms, TE =3.22 ms, flip angle = 7º, 192 sagittal slices, voxel size = 1 × 1 × 1 $mm^3$). RsfMRI data were obtained using an EPI sequence (TR = 2000 ms, TE =30 ms; flip angle = 90º; voxel size = 3.5 × 3.5 × 4.35

mm$^3$). Each resting-state scan consisted of 3 runs, with 8 min 6 sec per run. Written informed consent was obtained from each patient in accordance with guidelines and regulations previously approved by the IRB of China Rehabilitation Research Center.

#### Supplementary Note 2: Deep-learning algorithm descriptions

##### *FastSurferCNN*

FastSurferCNN is a deep learning architecture designed for anatomical segmentation of brain tissues, enabling computationally efficient segmentation of the entire brain into 95 distinct cortical and subcortical regions[4]. The FastSurferCNN employs a threefold approach, utilizing separate F-CNNs to process axial, coronal, and sagittal 2D slices. These three views are subsequently integrated into a comprehensive 3D segmentation at the view aggregation stage. Each F-CNN in FastSurferCNN comprises a sequence of dense encoder and decoder blocks, characterized by the incorporation of extensive skip connections that facilitate long-range information flow. Additionally, F-CNN integrates competitive dense blocks and maxout activations, and spatial information aggregation. These improvements serve to reduce the overall number of model parameters while simultaneously preserving the essential spatial information necessary for accurate segmentation of neuroanatomical structures, including cortical gray-matter regions. The final segmentation in FastSurferCNN is generated by aggregating probability maps from each of the three views through a weighted average mechanism. For further in-depth insights into the model architectures, implementations, and training procedures, please refer to the previous report[4].

##### *FastCSR*

FastCSR is a deep learning model designed for the time-consuming task of cortical surface reconstruction (CSR) through the utilization of implicit representations of cortical surfaces[5]. This model yields a remarkable speed enhancement, achieving an approximate processing speed 47 times faster than that of FreeSurfer in the context of CSR. FastCSR uses a 3D U-Net model to learn the level set, an implicit representation of the cortical surface. Within the level set representation, voxels with a zero value denote surface boundaries, while negative or positive values signify distances from these boundaries inward or outward, respectively. This level set is learning from the normalized T1w images and the white-matter segmentation outputs, following the FastSurferCNN processing. To predict the level set, we employ the state-of-the-art U-Net model, specifically the no-new-Net (nnU-Net) framework. Once the level set representations are learned, we proceed to derive the explicit triangular mesh representation of the white-matter cortical surface. This transformation is facilitated by a topology-preserving geometric deformable surface model[6]. Furthermore, we optimize the white-matter surfaces and generate pial cortical surfaces using a surface optimizing method implemented in FreeSurfer (see command *mris_make_surfaces*). More details on model architectures, implementations, and training could be found in the previous report and code repository (https://github.com/IndiLab/FastCSR) for a more comprehensive understanding of FastCSR[5].

***SUGAR***

Spherical Ultrafast Graph Attention framework for cortical surface Registration (SUGAR) is a deep-learning framework designed for both rigid and non-rigid cortical surface registration[7]. SUGAR exhibits a remarkable performance advantage, being approximately 12,000 times faster than the widely-used FreeSurfer in the context of cortical surface registration, when ensuring outperforming registration performance in to-target similarity and distortions. SUGAR's core architecture incorporates a U-Net-based spherical graph attention network (S-GAT). This network is trained at multiple resolutions using various standard subdivisions of an icosahedron. S-GAT relies on Euler angles to represent deformations and effectively leverages attention mechanisms to optimize performance. These mechanisms assign variable weights to different vertices, enhancing the precision of the registration process. Furthermore, SUGAR benefits from several strategic enhancements, including an optimized barycentric interpolation strategy, positional encoding strategy, and the integration of multiple loss functions that account for both similarity and distortions. Spherical data augmentation techniques are also employed, collectively resulting in improved computational efficiency and registration performance. For a comprehensive understanding of SUGAR, including intricate details related to model architectures, implementations, and training methodologies, we refer to our prior report[7] and the code repository (https://github.com/IndiLab/SUGAR).

***SynthMorph***

SynthMorph is a deep learning-based approach tailored for volumetric registration, also known as the spatial normalization[8], aiming to achieve robust, accurate, and computationally efficient alignment of previously unseen brain images. In its pursuit of high generalization, with a specific focus on achieving registration irrespective of image contrast variations, SynthMorph leverages a diverse set of synthetic data generated through a generative strategy. This approach exposes the neural networks to a wide spectrum of contrast variability, facilitating generalization across different imaging contrasts. SynthMorph adopts the U-Net architecture, a well-established framework employed in the VoxelMorph model[9]. Comprehensive details pertaining to model architectures, implementations, and training procedures can be referenced in a prior report[8] and a website page (https://martinos.org/malte/synthmorph/). In the DeepPrep framework, SynthMorph plays a pivotal role by independently executing affine and non-rigid registration processes. These procedures are applied to align native anatomical images with a standard volumetric template, such as the MNI152NLin6Asym template, yielding a deformation field. Subsequently, DeepPrep utilizes this deformation field in conjunction with boundary-based registration techniques to further process functional images.

**Supplementary Note 3: Minimum system requirements**

Operating system: Ubuntu 20.04 or higher

RAM + Swap space: at least 20 GB

Hard disk: at least 20 GB

*Optional but highly recommended:*

GPU VRAM: at least 12 GB

NVIDIA Driver: CUDA Toolkit 11.8 or higher

**Supplementary Tables**

**Supplementary Table.1 | software and toolbox dependencies**

| **Software used by DeepPrep** | | |
|---|---|---|
| *Processing step* | *Implementation* | *Software Version* |
| *Input data* | | |
| Data in NIfTI format | BIDS format | |
| *Anatomical processing* | | |
| brain extraction brain tissue segmentation | FastSufer | v1.1.0 |
| bias correction | SimpleITK's N4 | v2.1.1 |
| spatial normalization | SynthMorph | 1 |
| reconstruct cortical surfaces from T1w structural images | FastCSR | v1.0.0 |
| cortical surface registration | SUGAR | 1.0.0 |
| cortical surface parcellation | FreeSurfer | 7.2.0 |
| *BOLD processing* | | |
| BOLD reference image | niworkflows | 1.10.0 |
| head motion correction | FSL MCFLIRT | 6.0.5.1 |
| slice time correction | AFNI 3dTshift | 24.0.00 |
| susceptibility distortion correction | SDCFlow | 2.6.0 |
| coregistration | FreeSurfer bbregister | 7.2.0 |
| resampling EPI to standard space | SynthMorph | 1 |
| confounds estimation | In-house codes | |
| **Python packages used by DeepPrep** | | |
| *Packages* | | *Version* |
| fastcsr | | 1.0.0 |
| sugar | | 1.0.0 |
| nighres | | 1.5.0 |
| torch | | 2.0.1+cu118 |
| torchvision | | 0.15.2+cu118 |
| templateflow | | 23.1.0 |
| lapy | | 1.0.1 |
| voxelmorph | | 0.2 |
| tensorflow | | 2.11.1 |
| torch-geometric | | 2.2.0 |
| fmriprep | | 23.2.0 |
| mriqc | | 23.2.0 |
| nnunet | | 1.7.1 |
| dynaconf | | 3.2.3 |
| wand | | 0.6.11 |
| bids | | 0.0 |
| nipype | | 1.8.6 |
| niworkflows | | 1.10.0 |
| SimpleITK | | 2.3.0 |

| | | |
|---|---|---|
| sdcflows | | 2.8.1 |
| open3d | | 0.17.0 |
| onnxruntime | | 1.16.3 |
| pytorch3d | | 0.7.4 |
| python-redis-lock | | 4.0.0 |
| neurite | | 0.2 |

**Supplementary Table.2 | Characteristics of evaluation datasets**

| **No.** | **Dataset** | **# used subs** | **# total subs** | **age** | **source** | **state** | **scanner** |
|---|---|---|---|---|---|---|---|
| 1 | CoRR_HNU | 30 | 30 | 20-30 | http://fcon_1000.projects.nitrc.org/indi/CoRR/html/hnu_1.html | Healthy | GE MR750 3T |
| 2 | MSC | 10 | 10 | 24-36 | https://openneuro.org/datasets/ds000224/versions/1.0.3 | Healthy | Siemens TRIO 3T |
| 3 | UKB_20252_2.0 | 49300 | 49300 | 44-83 | https://biobank.ndph.ox.ac.uk/showcase/field.cgi?id=20252 | Healthy | Siemens Skyra 3T |
| 4 | UKB_20252_3.0 | 5215 | 5215 | 44-83 | https://biobank.ndph.ox.ac.uk/showcase/field.cgi?id=20252 | Healthy | Siemens Skyra |
| 5 | Mindboggle-101 | 97[a] | 101 | 19-61 | http://mindboggle.info/data | Healthy | Siemens, Philip |
| 6 | CRRC-Stroke | 19[b] | 218 | 37-71 | Collected in Beijing Bo'ai Hospital | Stroke | Philip Ingenia 3T |
| 7 | BTH-Glioma | 19[b] | 168 | 21-62 | Collected in Beijing Tiantan Hospital | Glioma | Siemens TrimTrio 3T |
| 8 | SHH-DoC | 15[b] | 38 | 31-53 | Collected in Shanghai Huashan Hospital | DoC | Siemens MAGNETOM 3T |

[a] Four cases have been excluded due to either poor data quality or the absence of necessary anatomical annotations.
[b] Cases were selected from the clinical datasets if FreeSurfer v6.0 failed to complete processing within 48 CPU hours.

**Supplementary Table.3 | Comparisons of anatomical segmentation in subcortical structures between fMRIPrep and DeepPrep**

| No. | Subcortical structures | fMRIPrep[a] | DeepPrep[b] | Contrast[c] |
|---|---|---|---|---|
| 1 | Left-Cerebral-White-Matter | 0.97±0.01 | 0.97±0.01 | 0.16% |
| 2 | Left-Lateral-Ventricle | 0.89±0.04 | 0.90±0.04 | 1.07% |
| 3 | Left-Inf-Lat-Vent | 0.67±0.14 | 0.68±0.15 | 1.59% |
| 4 | Left-Cerebellum-White-Matter | 0.84±0.07 | 0.85±0.08 | 1.40% |
| 5 | Left-Cerebellum-Cortex | 0.88±0.06 | 0.89±0.07 | 0.95% |
| 6 | Left-Thalamus-Proper | 0.90±0.02 | 0.91±0.02 | 0.18% |
| 7 | Left-Caudate | 0.88±0.03 | 0.89±0.02 | 0.16% |
| 8 | Left-Putamen | 0.85±0.03 | 0.86±0.03 | 0.90% |
| 9 | Left-Pallidum | 0.76±0.12 | 0.75±0.11 | -1.24% |
| 10 | 3rd-Ventricle | 0.85±0.07 | 0.87±0.07 | 2.09% |
| 11 | 4th-Ventricle | 0.85±0.04 | 0.87±0.07 | 1.85% |
| 12 | Brain-Stem | 0.92±0.01 | 0.93±0.04 | 1.42% |
| 13 | Left-Hippocampus | 0.88±0.04 | 0.88±0.04 | 0.27% |
| 14 | Left-Amygdala | 0.84±0.04 | 0.83±0.03 | -1.46% |
| 15 | CSF | 0.77±0.06 | 0.80±0.05 | 3.35% |
| 16 | Left-Accumbens-area | 0.69±0.09 | 0.69±0.07 | 0.13% |
| 17 | Left-VentralDC | 0.84±0.03 | 0.84±0.03 | 0.02% |
| 18 | Left-choroid-plexus | 0.42±0.07 | 0.49±0.07 | 15.40%* |
| 19 | Right-Cerebral-White-Matter | 0.97±0.01 | 0.97±0.01 | 0.13% |
| 20 | Right-Lateral-Ventricle | 0.89±0.05 | 0.90±0.05 | 1.58% |
| 21 | Right-Inf-Lat-Vent | 0.69±0.15 | 0.72±0.16 | 4.59% |
| 22 | Right-Cerebellum-White-Matter | 0.83±0.08 | 0.84±0.09 | 1.24% |
| 23 | Right-Cerebellum-Cortex | 0.88±0.06 | 0.89±0.07 | 0.78% |
| 24 | Right-Thalamus-Proper | 0.91±0.02 | 0.91±0.02 | 0.55% |
| 25 | Right-Caudate | 0.88±0.04 | 0.89±0.03 | 0.37% |
| 26 | Right-Putamen | 0.87±0.02 | 0.88±0.02 | 0.72% |
| 27 | Right-Pallidum | 0.77±0.07 | 0.78±0.07 | 0.54% |
| 28 | Right-Hippocampus | 0.88±0.03 | 0.89±0.03 | 0.53% |
| 29 | Right-Amygdala | 0.82±0.07 | 0.81±0.06 | -1.31% |
| 30 | Right-Accumbens-area | 0.70±0.06 | 0.68±0.04 | -2.61% |
| 31 | Right-VentralDC | 0.84±0.04 | 0.84±0.03 | 0.16% |
| 32 | Right-choroid-plexus | 0.45±0.07 | 0.51±0.06 | 11.80% |
| 33 | WM-hypointensities | 0.45±0.09 | 0.48±0.09 | 6.01% |
| 34 | CC_Posterior | 0.85±0.09 | 0.88±0.10 | 3.14% |
| 35 | CC_Mid_Posterior | 0.79±0.10 | 0.78±0.10 | -1.54% |
| 36 | CC_Central | 0.78±0.11 | 0.68±0.09 | -13.48%* |
| 37 | CC_Mid_Anterior | 0.80±0.13 | 0.70±0.10 | -12.29%* |
| 38 | CC_Anterior | 0.84±0.10 | 0.83±0.09 | -1.38% |

[a] Dice coefficients between automatic segmentations derived from fMRIPrep and manual segmentations
[b] Dice coefficients between automatic segmentations derived from DeepPrep and manual segmentations
[c] Percentages of differences in Dice coefficients derived from fMRIPrep and DeepPrep. Positive percentages indicate better performances of DeepPrep.
[*] two-tailed paired sample *t* tests, $p < 0.01$, FDR corrected

**Supplementary Table.4 | Comparisons of anatomical segmentation in cortical regions between fMRIPrep and DeepPrep**

| Cortical regions | left hemisphere | | | right hemisphere | | |
|---|---|---|---|---|---|---|
| | fMRIPrep[a] | DeepPrep[b] | contrast[c] | fMRIPrep[a] | DeepPrep[b] | contrast[c] |
| Caud. ant. cingulate | 0.68±0.15 | 0.67±0.15 | -1.13% | 0.76±0.14 | 0.79±0.15 | 3.86% |
| Caud. mid. frontal | 0.91±0.04 | 0.91±0.04 | 0.09% | 0.90±0.05 | 0.90±0.05 | 0.15% |
| Cuneus cortex | 0.79±0.04 | 0.79±0.04 | 0.11% | 0.82±0.06 | 0.82±0.06 | -0.04% |
| Entorhinal cortex | 0.76±0.08 | 0.74±0.10 | -3.35% | 0.76±0.10 | 0.71±0.13 | -5.99% |
| Fusiform gyrus | 0.82±0.05 | 0.82±0.05 | 0.07% | 0.85±0.06 | 0.85±0.06 | -0.25% |
| Inf. parietal | 0.85±0.05 | 0.85±0.05 | 0.12% | 0.85±0.05 | 0.85±0.05 | 0.13% |
| Inf. temporal | 0.82±0.06 | 0.82±0.06 | -0.02% | 0.86±0.04 | 0.86±0.04 | 0.04% |
| Isthmus | 0.77±0.06 | 0.82±0.05 | 5.81%* | 0.77±0.05 | 0.82±0.04 | 6.44%* |
| Lat. occipital | 0.87±0.04 | 0.87±0.04 | 0.10% | 0.86±0.05 | 0.86±0.05 | 0.02% |
| Lat. orbito frontal | 0.81±0.04 | 0.81±0.04 | -0.15% | 0.80±0.04 | 0.81±0.04 | 0.66% |
| Lingual gyrus | 0.92±0.03 | 0.92±0.03 | 0.54% | 0.92±0.03 | 0.93±0.03 | 0.33% |
| Med. orbito frontal | 0.78±0.07 | 0.78±0.07 | 0.85% | 0.70±0.08 | 0.72±0.08 | 2.56% |
| Mid. temporal | 0.80±0.04 | 0.80±0.05 | 0.02% | 0.83±0.04 | 0.83±0.04 | 0.13% |
| Parahippocampal | 0.78±0.03 | 0.85±0.03 | 8.22%* | 0.78±0.04 | 0.84±0.04 | 7.94%* |
| Paracentral lobule | 0.87±0.05 | 0.87±0.05 | 0.11% | 0.90±0.04 | 0.90±0.04 | 0.02% |
| Pars opercularis | 0.82±0.09 | 0.82±0.09 | -0.09% | 0.80±0.09 | 0.80±0.09 | 0.01% |
| Pars orbitalis | 0.44±0.11 | 0.44±0.11 | 1.02% | 0.53±0.12 | 0.54±0.12 | 0.13% |
| Pars triangularis | 0.73±0.10 | 0.73±0.10 | 0.42% | 0.77±0.09 | 0.77±0.10 | 0.04% |
| Pericalcarine | 0.88±0.06 | 0.89±0.06 | 0.84% | 0.89±0.05 | 0.90±0.05 | 0.61% |
| Postcentral | 0.89±0.05 | 0.89±0.05 | 0.05% | 0.90±0.06 | 0.90±0.06 | 0.03% |
| Posterior-cingulate | 0.85±0.07 | 0.86±0.07 | 1.02% | 0.89±0.06 | 0.90±0.06 | 1.11% |
| Precentral | 0.93±0.04 | 0.93±0.04 | -0.05% | 0.91±0.04 | 0.91±0.04 | -0.01% |
| Precuneus | 0.89±0.02 | 0.89±0.02 | 0.09% | 0.89±0.03 | 0.89±0.03 | 0.07% |
| Rost. ant. cingulate | 0.70±0.10 | 0.66±0.11 | -6.36% | 0.65±0.13 | 0.65±0.14 | 0.11% |
| Rost. mid. frontal | 0.79±0.05 | 0.79±0.05 | -0.01% | 0.78±0.06 | 0.78±0.06 | 0.04% |
| Sup. frontal | 0.80±0.05 | 0.80±0.05 | -0.05% | 0.80±0.05 | 0.80±0.05 | 0.04% |
| Sup. parietal | 0.84±0.04 | 0.84±0.04 | 0.08% | 0.84±0.05 | 0.84±0.05 | 0.06% |
| Sup. temporal | 0.83±0.03 | 0.83±0.03 | -0.08% | 0.83±0.04 | 0.83±0.04 | 0.00% |
| Supramarginal | 0.86±0.05 | 0.86±0.05 | 0.02% | 0.85±0.05 | 0.85±0.05 | 0.17% |
| Transverse temporal | 0.90±0.07 | 0.90±0.07 | 0.09% | 0.89±0.06 | 0.89±0.06 | 0.27% |
| Insula | 0.80±0.03 | 0.85±0.02 | 6.21%* | 0.84±0.03 | 0.84±0.03 | 0.81% |

[a] Dice coefficients between automatic segmentations derived from fMRIPrep and manual segmentations
[b] Dice coefficients between automatic segmentations derived from DeepPrep and manual segmentations
[c] Percentages of differences in Dice coefficients derived from fMRIPrep and DeepPrep. Positive percentages indicate better performances of DeepPrep.
[*] two-tailed paired sample *t* tests, $p < 0.01$, FDR corrected

#### Extended Figures

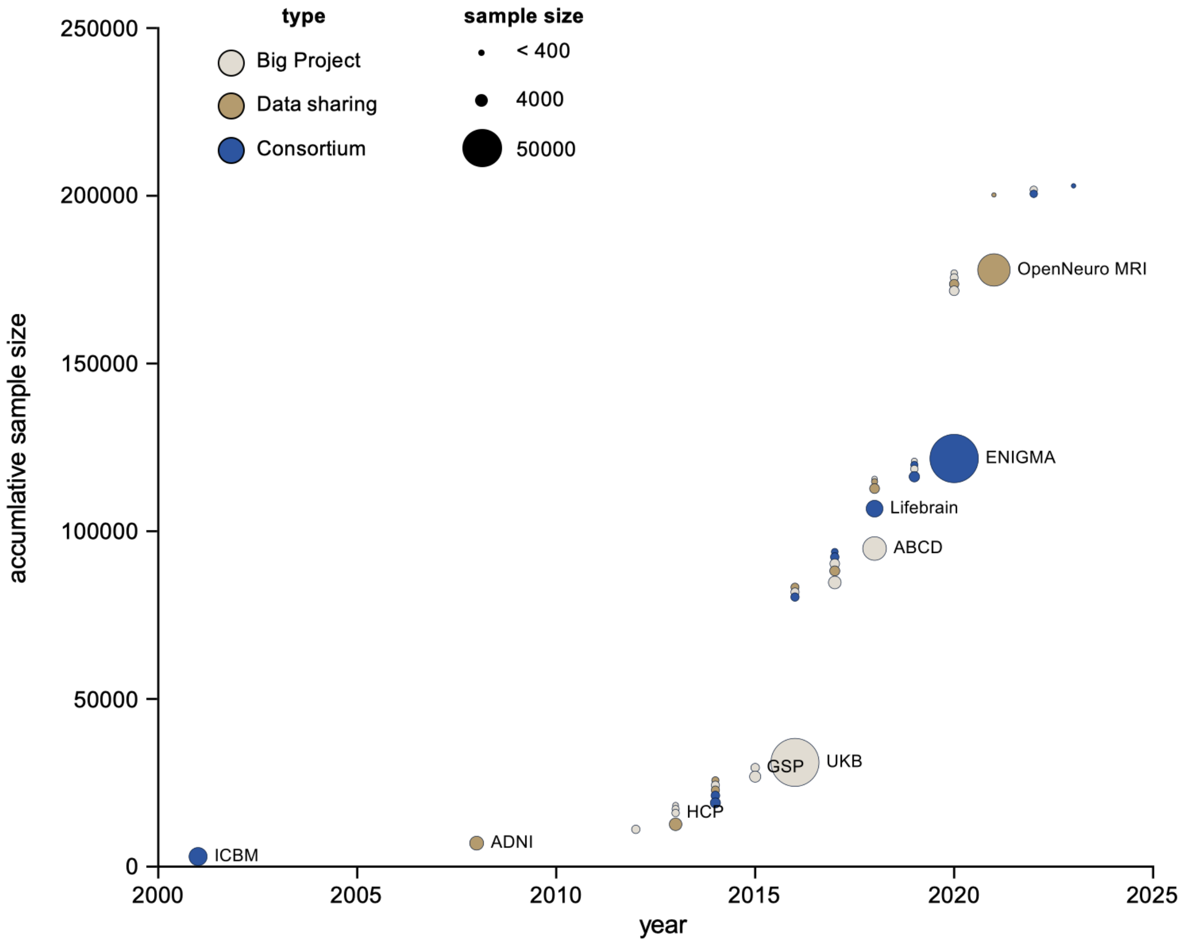


**Extended Data Fig.1 | Exponential growth of publicly available neuroimaging datasets from 2000 to 2023.** The figure depicts the cumulative sample size of publicly accessible neuroimaging datasets, which has exhibited exponential growth over the past two decades. A total of forty-two large-scale neuroimaging datasets were selected through a comprehensive meta-analysis spanning from the year 2000 to 2023. These datasets are categorized into three representative types: 'big project' (white circles), supported by a single funding source, 'consortium' (blue circles), resulting from international collaborations involving multiple funding sources, and 'sharing data platform' (sand circles), such as the openneuro, which serves as a data repository. The size of each circle in the plot corresponds to the sample size of the respective dataset. The project names of some representative datasets are labeled next to the corresponding circles in the dataset. Stacked circles represent cases where multiple different datasets were released in a single year. The Y-axis represents the cumulative sample size, demonstrating a pronounced exponential increase over time. This trend signifies the rapid transition of the neuroimaging field into the era of big data.

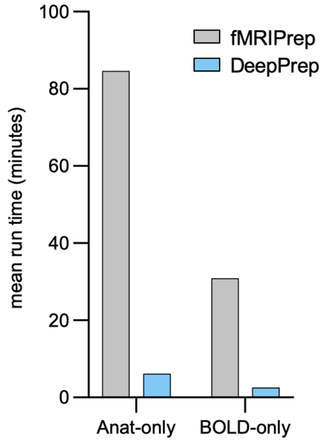


**Extended Data Fig. 2 | DeepPrep shows over 10-fold accelerations in preprocessing both anatomical and functional images.** In addition to the comparison of processing times for the entire pipeline between fMRIPrep and DeepPrep, we performed assessments focusing on the separate processing times for anatomical and functional images. When considering anatomical preprocessing with the '--anat-only' flag, DeepPrep completed the task in an impressive 6.15 minutes, marking a substantial 13.8-fold increase in speed compared to fMRIPrep, which took 84.66 minutes. Similarly, when processing functional images with the '--bold-only' flag, DeepPrep exhibited significant efficiency, finishing the task in 2.55 minutes, a 12.09-fold improvement over fMRIPrep, which required 30.84 minutes. During functional image processing, preprocessed anatomical images were provided at the outset.

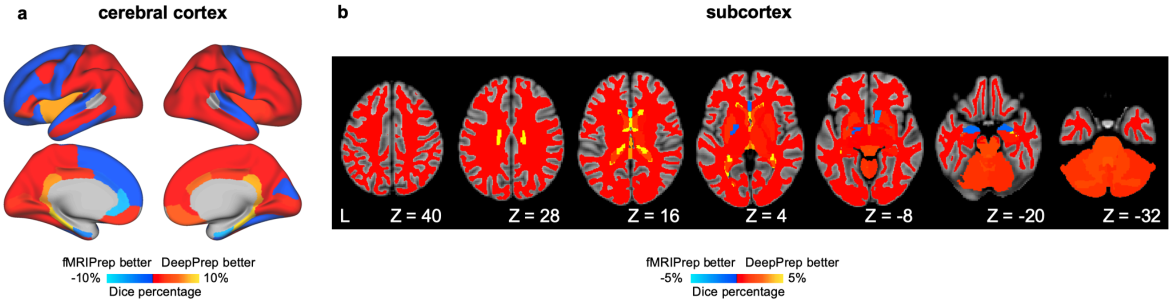


**Extended Data Fig. 3 | Comparable performance in brain tissue segmentation of DeepPrep and fMRIPrep.** DeepPrep's performance in brain tissue segmentation was assessed and compared with that of fMRIPrep. To demonstrate segmentation accuracy, both DeepPrep and fMRIPrep were applied to the

Mindboggle-101 dataset. This dataset includes anatomical images and brain regions manually segmented 'ground truth' data by experts. The accuracy of automatic segmentations produced by these pipelines was evaluated by calculating Dice coefficients, which measure the similarity of brain regions between the automatic segmentations and the ground truth data. Comparison of segmentation accuracy between DeepPrep and fMRIPrep was conducted by evaluating the percentage difference in Dice coefficients for each brain region. Positive percentages indicate higher accuracy achieved by DeepPrep, while negative percentages indicate higher accuracy for fMRIPrep. **a)** 47 out of 62 cortical regions displays positive percentages. **b)** Similar comparisons were performed for 39 subcortical regions, where 31 regions show positive percentages, suggesting better performance of DeepPrep in cortical anatomical parcellation compared to fMRIPrep. All statistics are reported in Supplementary Table 3 and Table 4.

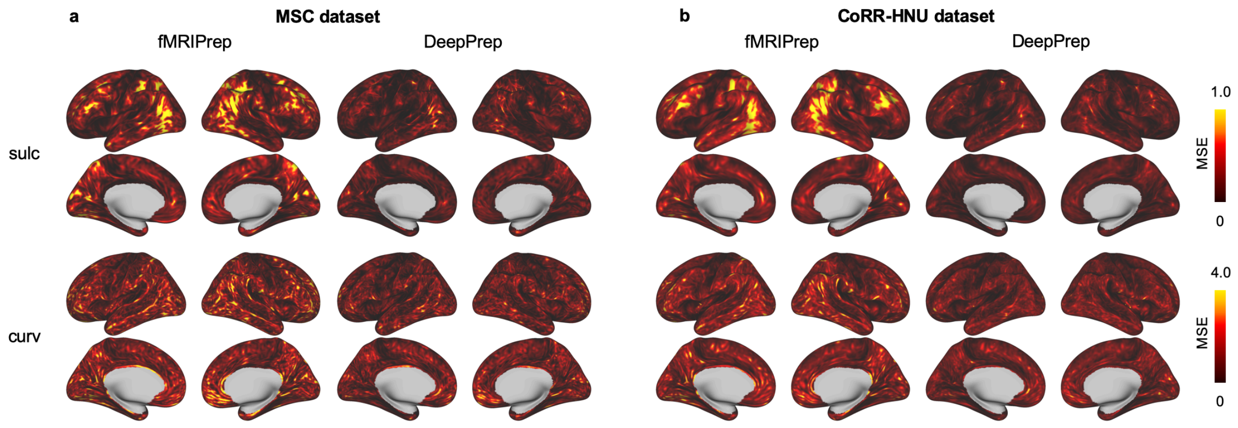


**Extended Data Fig. 4 | Improved similarity of cortical morphometric patterns with target atlases by DeepPrep.** Preprocessing pipelines reconstructed individual cortical surfaces and aligned surface morphometrics with target atlases. Both fMRIPrep and DeepPrep were used to process anatomical images from two datasets: (a) the MSC dataset and (b) the CoRR-HNU dataset. To evaluate the performance in surface reconstruction and registration, we conducted a comparison using three morphometric measures: sulcal depth (top row), curvature (bottom row). The assessment was based on the mean square error (MSE) when contrasting these measures with the target atlases. Across all morphometric measures and datasets, DeepPrep consistently shows lower MSE values than fMRIPrep.

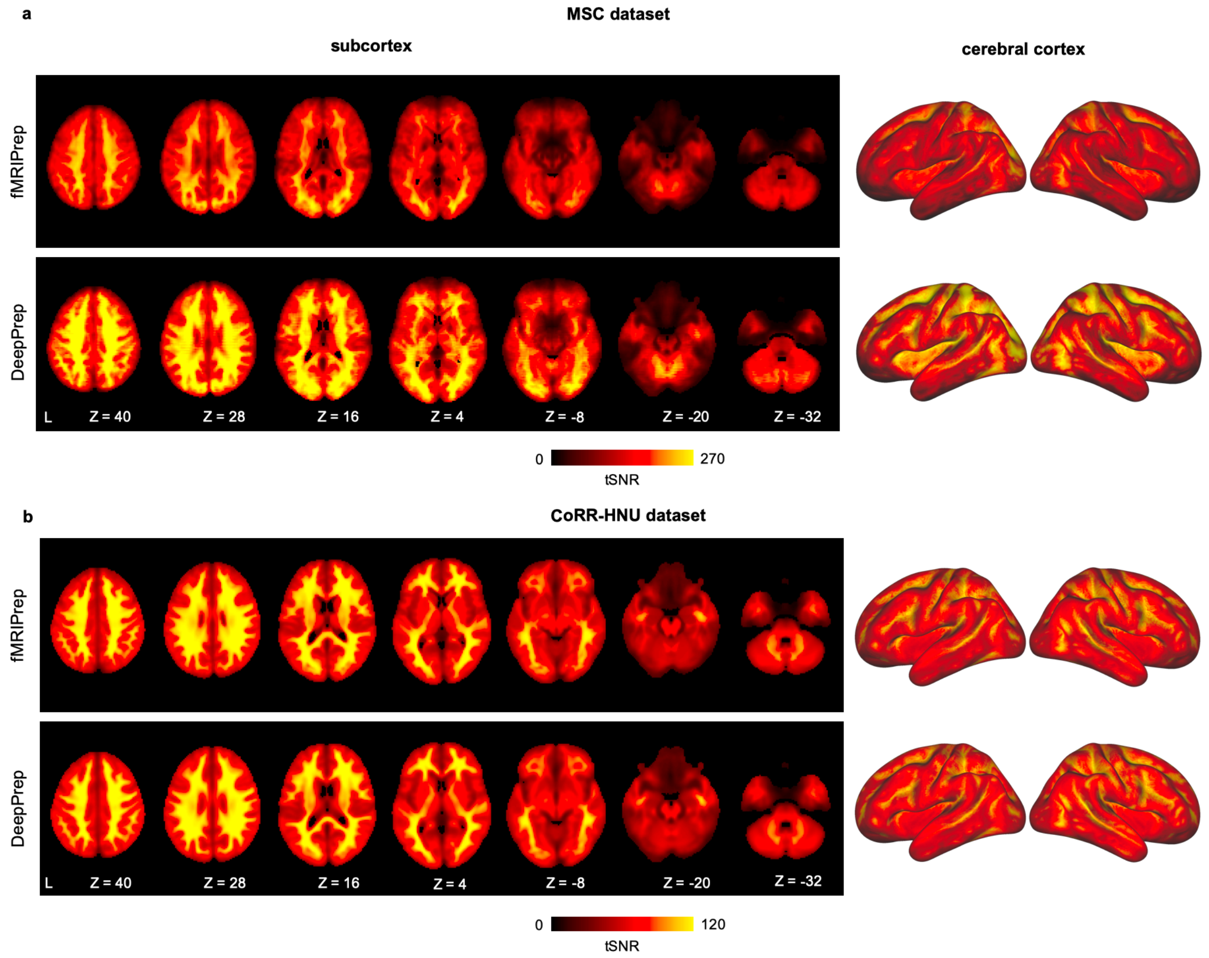


**Extended Data Fig. 5 | Enhanced temporal signal-to-noise ratio with DeepPrep.** Following the spatial normalization of BOLD-fMRI signals to the MNI152NLin6Asym volumetric template, we calculated the temporal signal-to-noise ratio (tSNR) for the BOLD-fMRI signals. DeepPrep consistently exhibited higher tSNR values in whole brains (left) and cortical surfaces (right, fasverage6) compared to fMRIPrep in both the **(a)** MSC and **(b)** CoRR-HNU datasets.

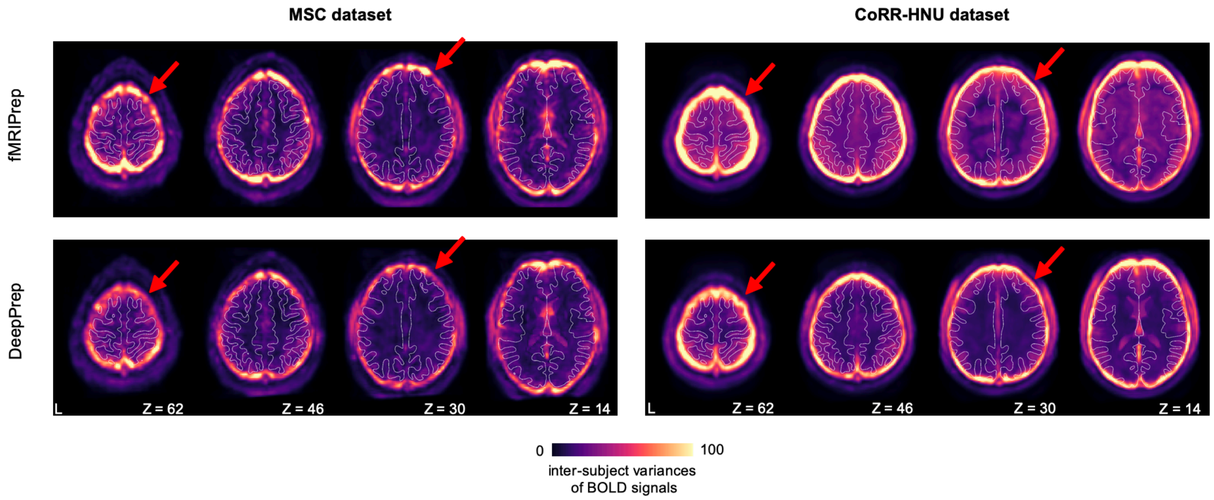


**Extended Data Fig. 6 | Reduced inter-subject variance in average BOLD signals with DeepPrep.** Following spatial normalization, we assessed the voxel-wise inter-subject variances in average BOLD signals across subjects from the MSC and CoRR-HNU datasets. Lighter colors in the figures indicate larger inter-subject variances. The white matter boundaries are delineated in white on the MNI152NLin6Asym volumetric template. While both datasets and preprocessing pipelines exhibited border effects, DeepPrep consistently demonstrated smaller inter-subject variance compared to fMRIPrep, as highlighted by red arrows in representative brain regions. The reduced inter-subject variances observed with DeepPrep suggest its robust performance in spatial normalization.

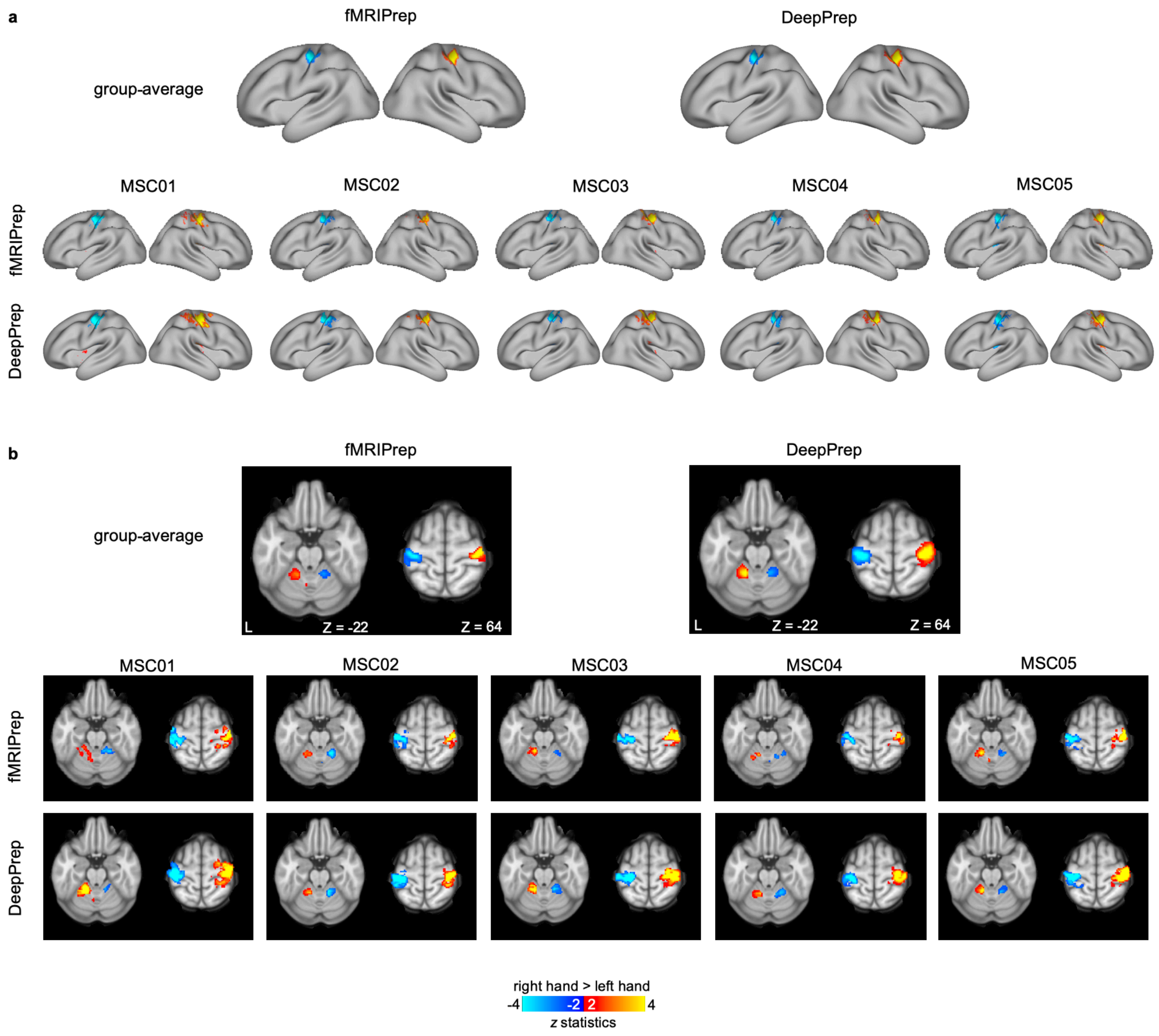


**Extended Data Fig. 7 | Comparable patterns and strength of task-evoked responses between DeepPrep and fMRIPrep.** DeepPrep and fMRIPrep were employed for preprocessing the motor task-evoked fMRI in the MSC dataset. Subsequent processing steps, including high-pass filtering and general linear modeling, were consistent between the two pipelines. (a) Contrast maps derived from the surface-based pipeline depicting the activation patterns for the right hand vs. right hand task are presented on a cortical surface template. (b) Volumetric pipelines yielded volumetric contrast maps, shown in a volumetric template. Both in group-average and individual contrast maps, the patterns and strengths of task-evoked responses are comparable between DeepPrep and fMRIPrep.

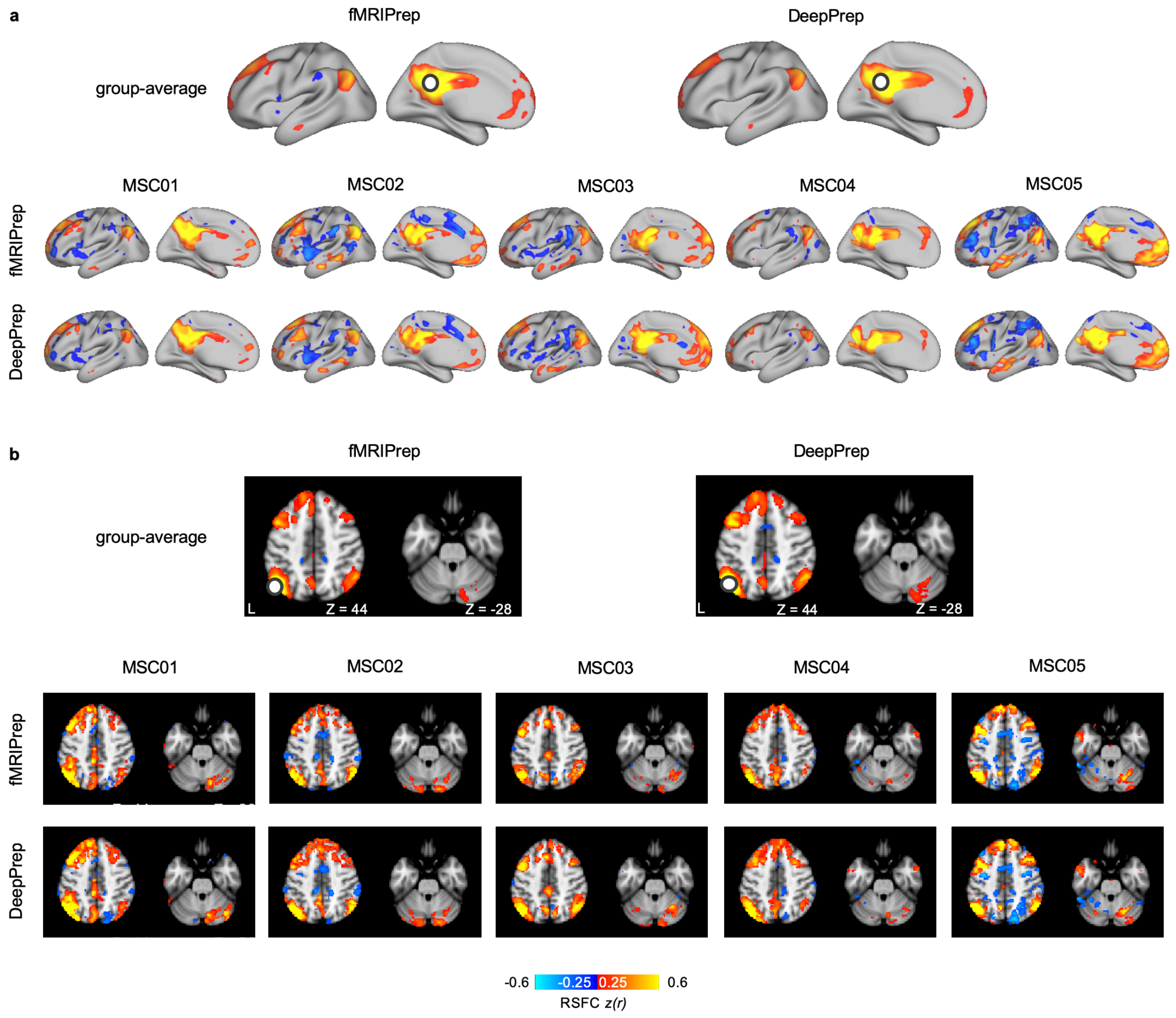


**Extended Data Fig. 8 | Comparable seed-based functional connectivity between DeepPrep and fMRIPrep.** DeepPrep and fMRIPrep were utilized to preprocess resting-state fMRI data from the MSC dataset. Subsequent processing steps, including band-pass filtering (0.01-0.08 Hz) and confound regression, were consistent between the two pipelines. Functional connectivity maps, derived from a 6-mm spherical seed ROI placed in the left posterior cingulate cortex (MNI coordinate = -2, -53, 26) and left angular gyrus (MNI coordinate = -49, -63, 45), are presented by the white circles. The seed-based functional connectivity maps derived from surface and volumetric pipelines are displayed within (a) a cortical surface template and (b) a volumetric template, respectively. In both group-average and individual connectivity maps, the patterns and strengths of functional connectivity are comparable between DeepPrep and fMRIPrep. This observation highlights the similarity in their performance in analyzing seed-based functional connectivity.

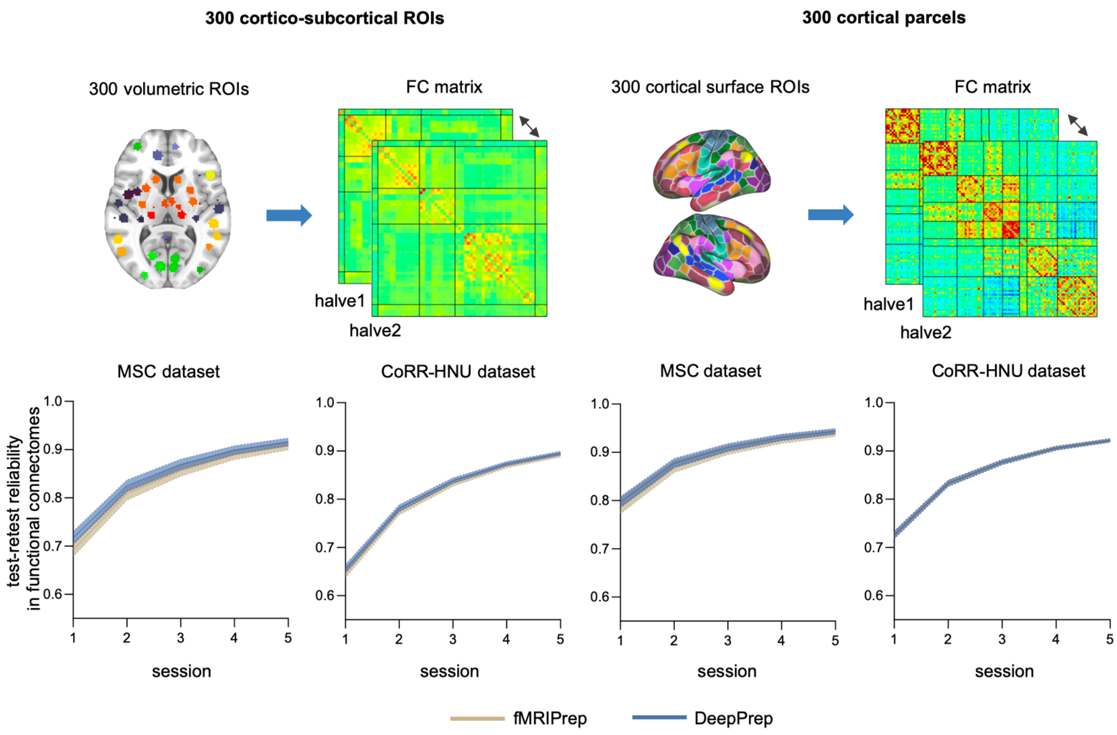


**Extended Data Fig. 9 | Comparable test-retest reliability of functional connectomes between DeepPrep and fMRIPrep.** To estimate whole-brain and cortical functional connectomes, we employed 300 volumetric ROIs encompassing both cortical and subcortical regions, as well as Schaefer's 300 cortical surface ROIs, respectively. Test-retest reliability was quantified by measuring the similarity between functional connectomes derived from two independent segments randomly sampled from one or multiple sessions in the repeated measured datasets, namely, the MSC dataset and the CoRR-HNU dataset. To elucidate the relationship between test-retest reliability and data amounts, we assessed reliability using one to five sessions. DeepPrep and fMRIPrep consistently exhibited comparable test-retest reliability in functional connectomes, with no significant differences observed (two-tailed paired-sample t-tests, FDR-corrected p-values > 0.05). The shaded regions in the curves represent standard deviations.

### Supplementary Figures

#### Supplementary Fig. 1 | An example of output files

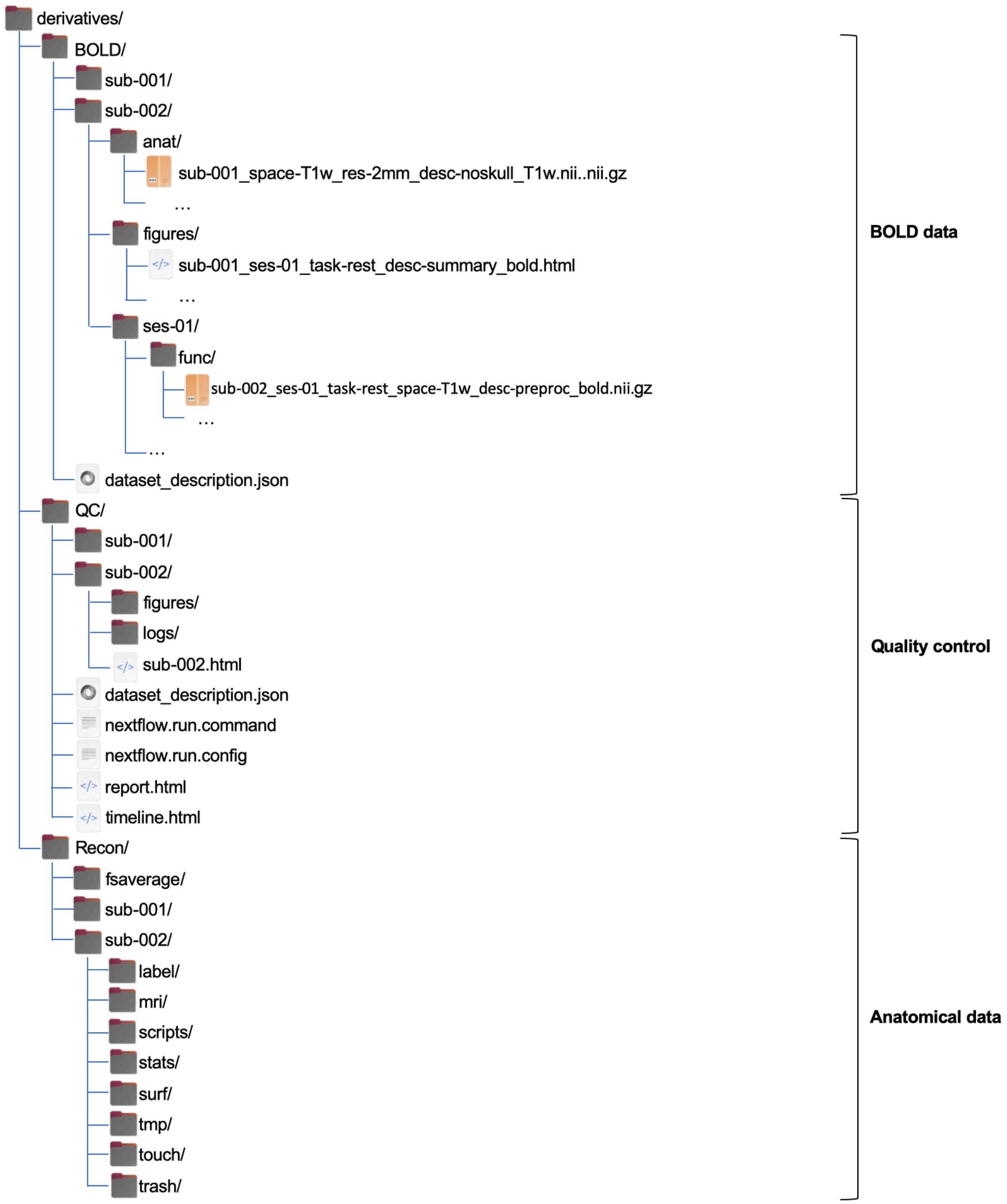

### Supplementary Fig. 2 | An example of the DeepPrep report

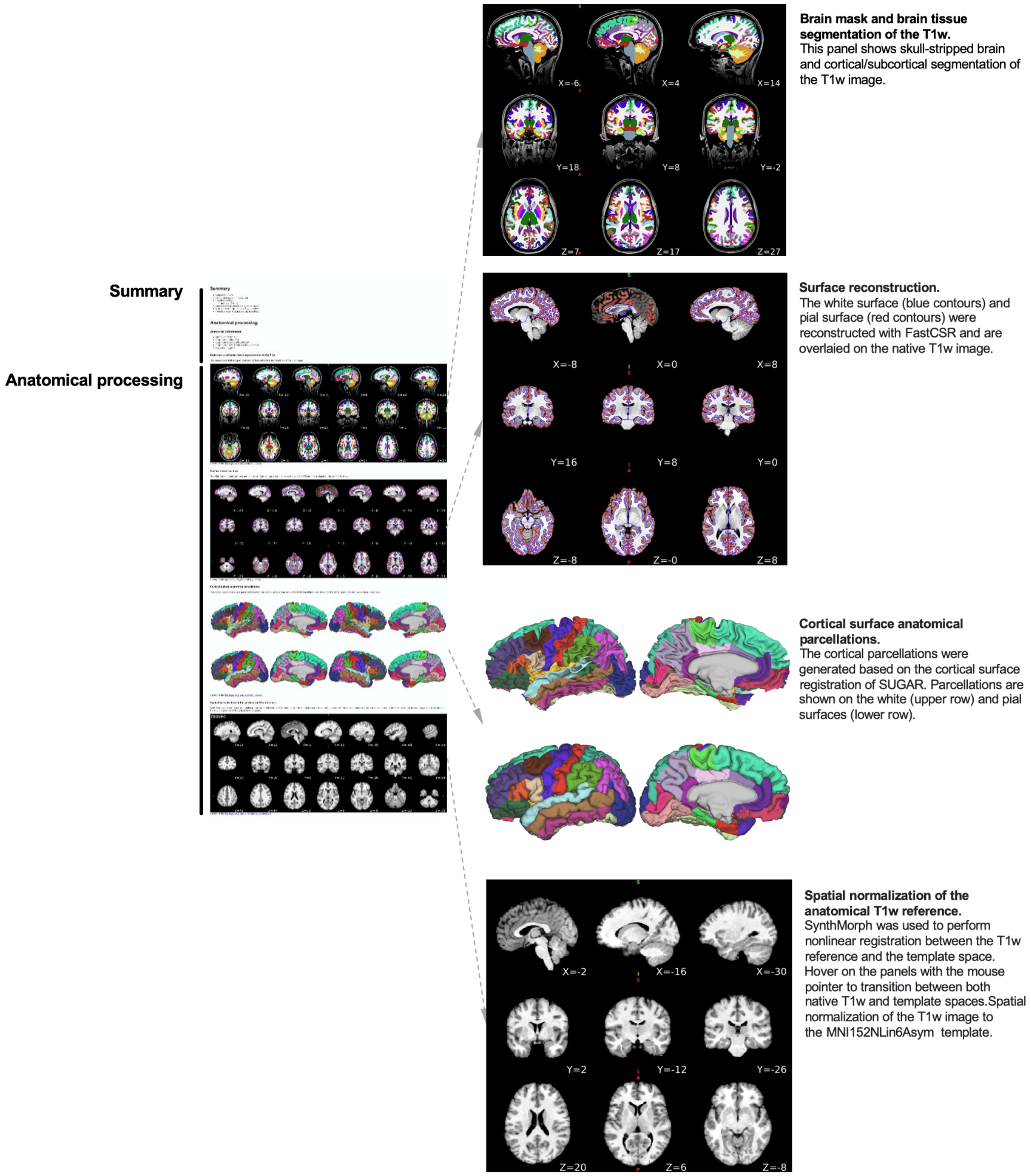

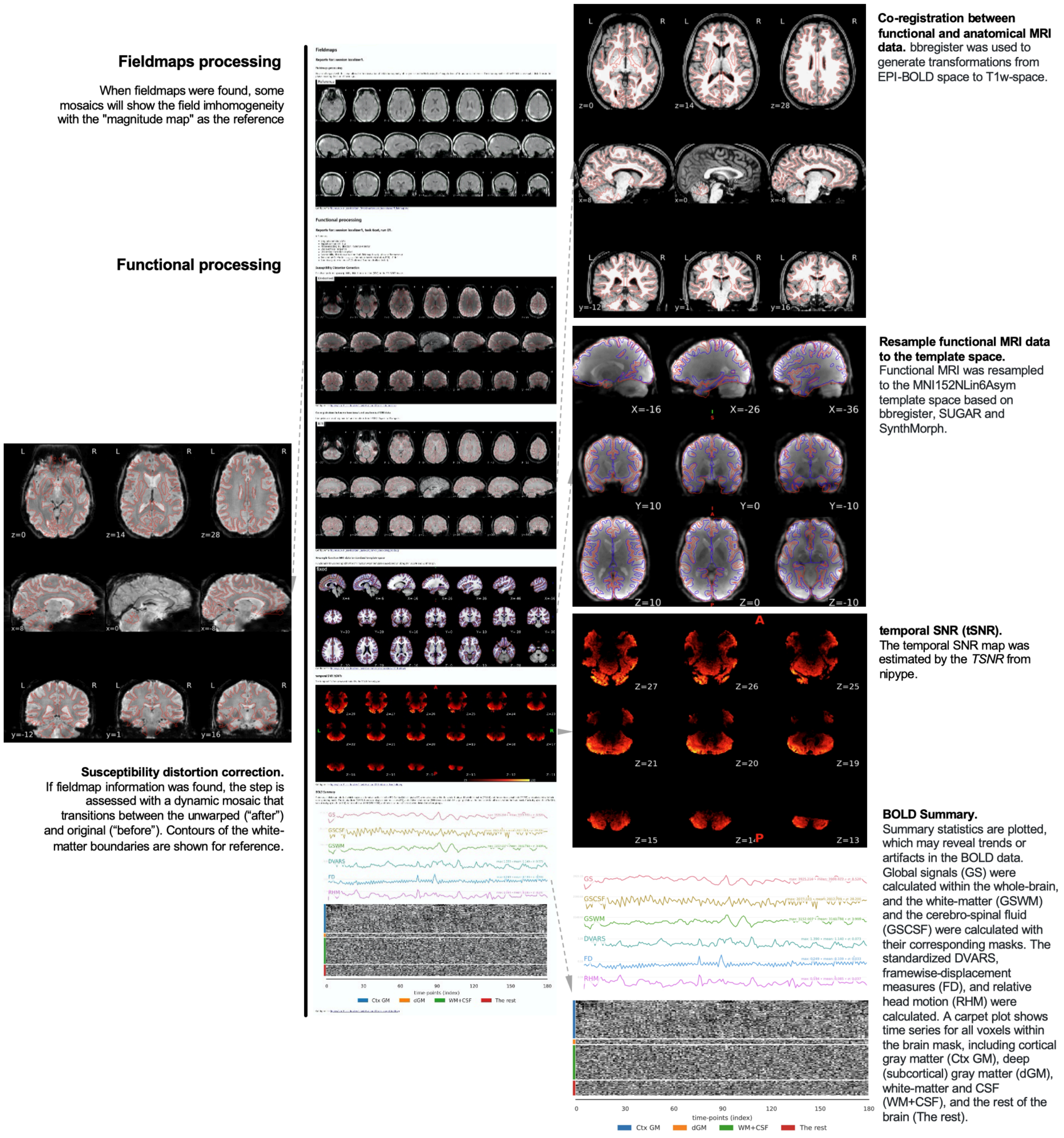
Fieldmaps processing
When fieldmaps were found, some mosaics will show the field imhomogeneity with the "magnitude map" as the reference
Functional processing
Susceptibility distortion correction.
If fieldmap information was found, the step is assessed with a dynamic mosaic that transitions between the unwarped ("after") and original ("before"). Contours of the white-matter boundaries are shown for reference.
Co-registration between functional and anatomical MRI data. bbregister was used to generate transformations from EPI-BOLD space to T1w-space.
Resample functional MRI data to the template space.
Functional MRI was resampled to the MNI152NLin6Asym template space based on bbregister, SUGAR and SynthMorph.
temporal SNR (tSNR).
The temporal SNR map was estimated by the TSNR from nipype.
BOLD Summary.
Summary statistics are plotted, which may reveal trends or artifacts in the BOLD data. Global signals (GS) were calculated within the whole-brain, and the white-matter (GSWM) and the cerebro-spinal fluid (GSCSF) were calculated with their corresponding masks. The standardized DVARS, framewise-displacement measures (FD), and relative head motion (RHM) were calculated. A carpet plot shows time series for all voxels within the brain mask, including cortical gray matter (Ctx GM), deep (subcortical) gray matter (dGM), white-matter and CSF (WM+CSF), and the rest of the brain (The rest).
GS
GSCSF
GSWM
DVARS
FD
RHM
time-points (index)
Ctx GM
dGM
WM+CSF
The rest

**Supplementary Fig. 3 | An example of the runtime report**

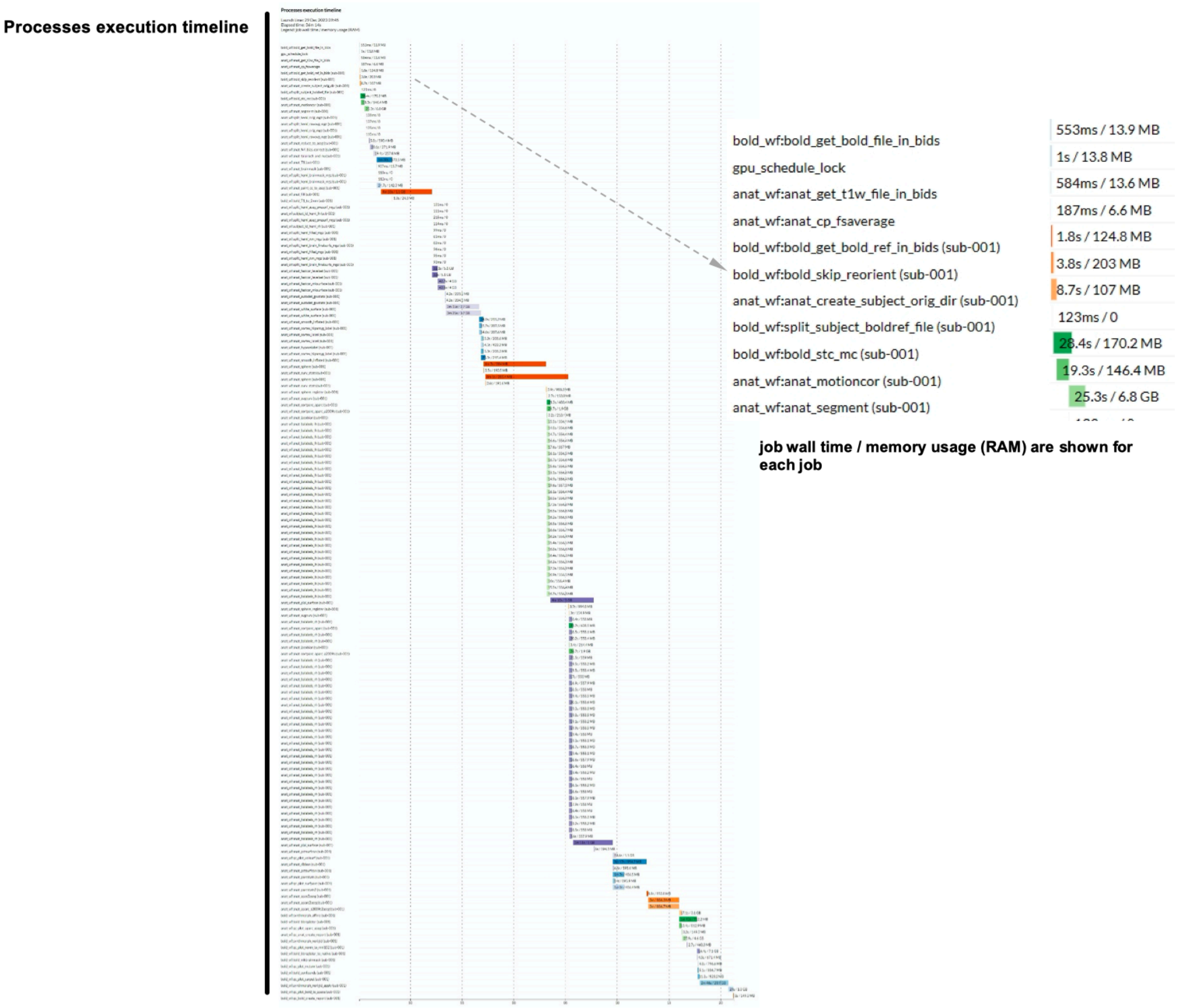
